# Characterization of the Human Immunodeficiency Virus (HIV-1) Envelope Glycoprotein Conformational States on Infectious Virus Particles

**DOI:** 10.1101/2022.12.01.518799

**Authors:** Hanh T. Nguyen, Qian Wang, Saumya Anang, Joseph G. Sodroski

## Abstract

Human immunodeficiency virus (HIV-1) entry into cells involves triggering of the viral envelope glycoprotein (Env) trimer ((gp120/gp41)_3_) by the primary receptor, CD4, and coreceptors, CCR5 or CXCR4. The pretriggered (State-1) conformation of the mature (cleaved) Env is targeted by broadly neutralizing antibodies (bNAbs), which are inefficiently elicited compared with poorly neutralizing antibodies (pNAbs). Here we characterize variants of the moderately triggerable HIV-1_AD8_ Env on virions produced by an infectious molecular proviral clone; such virions contain more cleaved Env than pseudotyped viruses. We identified three types of cleaved wild-type AD8 Env trimers on virions: 1) State-1-like trimers preferentially recognized by bNAbs and exhibiting strong subunit association; 2) trimers recognized by pNAbs directed against the gp120 coreceptor-binding region and exhibiting weak, detergent-sensitive subunit association; and 3) a minor gp41-only population. The first Env population was enriched and the other Env populations reduced by introducing State-1-stabilizing changes in the AD8 Env or by treatment of the virions with crosslinker or the State-1-preferring entry inhibitor, BMS-806. These stabilized AD8 Envs were also more resistant to gp120 shedding induced by a CD4-mimetic compound or by incubation on ice. Conversely, a State-1-destabilized, CD4-independent AD8 Env variant exhibited weaker bNAb recognition and stronger pNAb recognition. Similar relationships between Env triggerability and antigenicity/shedding propensity on virions were observed for other HIV-1 strains. Our results show that State-1 Envs on virions can be significantly enriched by optimizing Env cleavage; stabilizing the pretriggered conformation by Env modification, crosslinking or BMS-806 treatment; strengthening Env subunit interactions; and using CD4-negative producer cells.

**IMPORTANCE:** Efforts to develop an effective HIV-1 vaccine have been frustrated by the inability to elicit broad neutralizing antibodies that recognize multiple virus strains. Such antibodies are able to bind a particular shape of the HIV-1 envelope glycoprotein trimer, as it exists on a viral membrane but before engaging receptors on the host cell. Here, we establish simple yet powerful assays to characterize the envelope glycoproteins in a natural context on virus particles. We find that, depending on the HIV-1 strain, some envelope glycoproteins change shape and fall apart, creating decoys that can potentially divert the host immune response. We identify requirements to keep the relevant envelope glycoprotein target for broad neutralizing antibodies intact on virus-like particles. These studies suggest strategies that should facilitate efforts to produce and use virus-like particles as vaccine immunogens.

## INTRODUCTION

Human immunodeficiency virus type 1 (HIV-1) entry into target cells is mediated by the viral envelope glycoprotein (Env) trimer, which is composed of three gp120 exterior subunits and three gp41 transmembrane subunits (1, 2). In infected cells, Env is synthesized as an uncleaved precursor in the rough endoplasmic reticulum (ER), where signal peptide cleavage, folding, trimerization, and the addition of high-mannose glycans take place (3–6). Exiting the ER, the trimeric gp160 Env precursor follows two pathways to the cell surface (7). In the conventional secretory pathway, the Env precursor transits through the Golgi compartment, where it is cleaved into gp120 and gp41 subunits and is further modified by the addition of complex sugars (8–11). These mature Envs are transported to the cell surface and are incorporated into virions (7). In the second pathway, the gp160 precursor bypasses the Golgi compartment and traffics directly to the cell surface; these uncleaved gp160 Envs lack complex carbohydrates and are excluded from virions (7).

Single-molecule fluorescence resonance energy transfer (smFRET) experiments indicate that, on virus particles, the Env trimer exists in three conformational states (States 1 to 3) (12). From its pretriggered conformation (State 1), the metastable Env trimer interacts with the receptors, CD4 and CCR5 or CXCR4, and undergoes transitions to lower-energy states (13–16). Initially, the engagement with CD4 induces an asymmetric intermediate Env conformation, with the CD4-bound protomer in State 3 and the unliganded protomers in State 2 (17). Binding of additional CD4 molecules to the Env trimer then induces the full CD4-bound, prehairpin intermediate conformation, with all three Env protomers in State 3 (12, 17). An extended coiled coil consisting of the heptad repeat (HR1) region of gp41 is exposed in the prehairpin intermediate (18–21). State-3 Env protomers subsequently interact with CCR5 or CXCR4 coreceptors to trigger the formation of a gp41 six-helix bundle, a process that results in fusion of the viral and target cell membranes (22–26).

Env is the only virus-specific molecule exposed on the viral surface and thus represents the major target for host neutralizing antibodies (27–29). Env strain variability, heavy glycosylation, conformational flexibility and structural heterogeneity are thought to contribute to HIV-1 persistence by diminishing the elicitation and binding of neutralizing antibodies (27–32). During natural infection, high titers of antibodies are elicited that recognize the gp160 Env precursor, which samples multiple conformations, and disassembled Envs (shed gp120, gp41 six-helix bundles) (33–45). These antibodies are poorly neutralizing because they fail to recognize the mature functional Env trimer, which mainly resides in State 1 (12,36,38,41–49). After years of infection, some HIV-1-infected individuals generate broadly neutralizing antibodies (bNAbs), most of which recognize the pretriggered (State-1) Env conformation (12,46–58). Passively administered monoclonal bNAbs are protective in animal models of HIV-1 infection, suggesting that the elicitation of bNAbs is an important goal for vaccines (59–63). Unfortunately, bNAbs have not been efficiently and consistently elicited in animals immunized with current vaccine candidates, including stabilized soluble gp140 (sgp140) SOSIP.664 trimers (64–75). Soluble Env trimers often elicit strong strain-restricted neutralizing antibodies or poorly neutralizing antibodies (pNAbs) targeting the gp120 V3 variable loop, neo-epitopes at the base of the trimer, or holes in the glycan shield (68,76–84).

Differences in the antigenicity, glycosylation and conformation of sgp140 SOSIP.664 trimers and the pretriggered (State-1) membrane Env have been observed (11,85–96). Given the requirement for bNAbs to recognize conserved and conformationally-specific elements on the pretriggered (State-1) Env (12,46–49), even small differences from the native State-1 Env might affect immunogen efficacy. First, differences in the composition of the glycan shield of sgp140 SOSIP.664 trimers and membrane Env (11,90–93) could hamper the elicitation of bNAbs, most of which recognize epitopes that include glycan components or are surrounded by glycans that influence antibody access (28,29,32,97–101). Currently, the only way to obtain a glycan shield resembling that of the virion Env spike is to produce the Env trimer immunogen in a membrane-anchored form. Second, membrane Env trimer immunogens also have the capacity to present the full set of quaternary bNAb epitopes (including the gp41 membrane-proximal external region (MPER)) to the immune system. Although the MPER is important for the maintenance of the State-1 Env conformation (102–113), the MPER is removed from sgp140 SOSIP.664 trimers to prevent aggregation (32,114-117). Finally, immunodominant responses against neo-epitopes artefactually created in the base of soluble timers (79, 84) could be eliminated by using membrane Env trimer immunogens. Overall, these considerations indicate that a membrane Env immunogen may have considerable advantages in presenting a native State-1 Env conformation to the host immune system.

Virus-like particles (VLPs) are an attractive platform for HIV-1 membrane Env immunogens, as virions represent a natural environment for the functional Env trimer and should accurately reproduce the relevant bNAb target. The relative enrichment of Golgi-passaged Env in virions and VLPs results in Env trimers that are cleaved, authentically glycosylated and imbedded in a cholesterol/sphingomyelin-rich (lipid raft-like) membrane, all of which are conducive to the maintenance of a pretriggered (State-1) conformation (40-45,107-113,118). Although the low Env content of VLPs (mimicking the natural 10-14 Env spikes per virion (119, 120)) is not an absolute barrier to their use as immunogens, methods for efficient VLP production must be optimized. Of potentially greater importance is the quality of the Env on VLPs; indeed, several studies have documented significant Env heterogeneity in VLP preparations (121–137). Proteolytically mature Env is enriched in virions produced by infected cells; however, systems overexpressing HIV-1 Gag and Env often produce VLPs with significant levels of uncleaved Env (7,124–127,134,137). Uncleaved Env, which is flexible and prone to assume non-State-1 conformations recognizable by pNAbs (40–45), is a highly undesirable contaminant in any immunogen attempting to focus the host antibody response on the pretriggered (State-1) Env conformation. Envs with disrupted subunit interactions, manifest in the extreme case by shedding of gp120 from the trimer, represent other potential sources of heterogeneity (137–139).

Here, we study Env conformation on infectious HIV-1 particles, examining both cleaved and uncleaved Env populations. We compare viral pseudotypes and virions produced by infectious molecular proviral clones (IMCs). We evaluate the antigenicity and propensity to shed gp120 for Envs from different HIV-1 strains and Envs modified to stabilize distinct conformational states. These studies provide useful assays and reagents for the study of Env in a natural membrane context and can guide the optimization of Env quality in VLPs.

## RESULTS

### Comparison of systems transiently expressing virus particles

To identify a virus-producing system that could yield sufficient levels of cleaved Env for detailed analysis, we compared 293T cells transiently expressing either pseudotyped viruses or virions produced by an infectious molecular proviral clone (IMC). Viruses pseudotyped by the tier-2 primary HIV-1_AD8_ Env were produced by cotransfection of a plasmid expressing the AD8 Env and an *env*-negative provirus vector, pNL4-3.ΔEnv (originally pNL4-3.Luc.R-E-from the NIH HIV Reagent Program). In the pNL4-3.AD8 IMC, the NL4-3 *env* gene was replaced by that of AD8. The AD8 Envs produced by both expression systems have a signal peptide and C-terminal gp41 cytoplasmic tail from HXBc2/NL4-3 Envs. As seen in Fig. 1A, pseudotyped virus particles contained mostly uncleaved Env with little cleaved Env. By contrast, virions produced by the IMC incorporated at least as much Env per particle, but with ∼5.7-fold more gp120 relative to gp160. The pNL4-3.AD8 IMC also produced virions with efficiently cleaved Env in HeLa cells (Fig. 1A). Varying the transfected amounts of the Env-expressing and *env*-negative proviral plasmids failed to increase the gp120:gp160 ratio of the pseudovirus particles to a level comparable to that of the IMC-produced virions (Fig. 1B). These observations indicated that the IMC system could produce virus particles with sufficient quantities of cleaved Env for our study.

**FIG 1.**
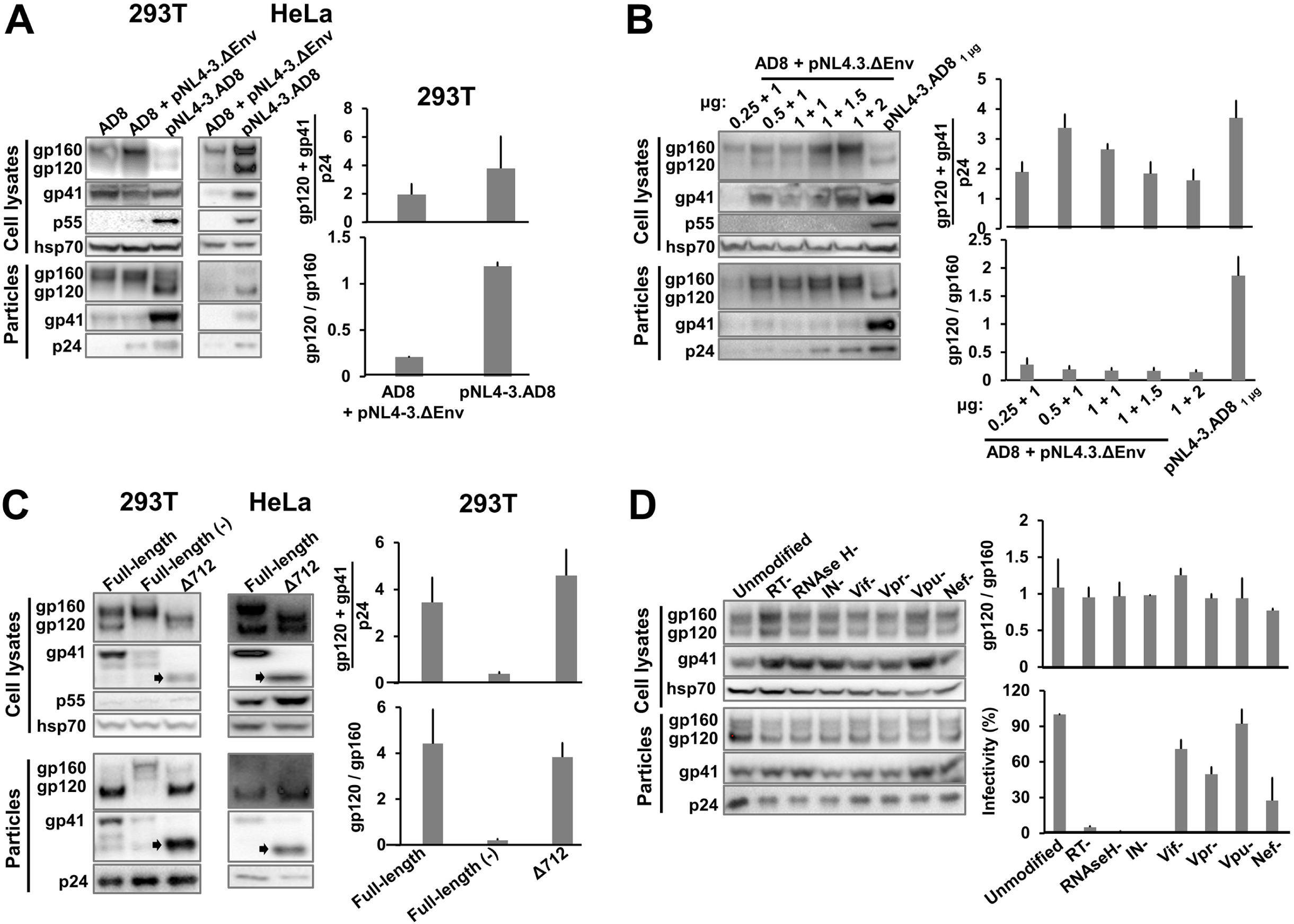
Comparison of virus pseudotypes and virions produced by an infectious molecular clone (IMC). (A) 293T cells and HeLa cells were transfected with the pSVIIIenv AD8 plasmid expressing the full-length HIV-1_AD8_ Env alone (AD8) or together with the pNL4-3.ΔEnv plasmid, or the pNL4-3.AD8 IMC alone (2 μg of each plasmid was used, whether alone or in combination). When the pSVIIIenv AD8 expressor plasmid was used, a plasmid expressing the HIV-1 Tat protein was also transfected at an 8:1 Env:Tat weight ratio. Forty-eight to seventy-two hours later, the cell supernatants were collected, filtered through a 0.45-μm membrane and centrifuged at 14,000 x g for 1 h at 4°C. In parallel, the cells were lysed. Precipitated particles and clarified cell lysates were Western blotted with a goat anti-gp120 antibody, the 4E10 anti-gp41 antibody, the rabbit anti-hsp70 antibody and the mouse anti-p24 serum. (B) 293T cells were transfected with the pSVIIIenv AD8 plasmid expressing the full-length HIV-1_AD8_ Env and the pNL4-3.ΔEnv plasmid at indicated weight ratios, or the pNL4-3.AD8 IMC alone. Cell lysates and virus particles were subsequently prepared and Western blotted as described above. (C) 293T cells and HeLa cells were transfected with the IMCs expressing the full-length AD8 Env, full-length Env with a defective cleavage site (−) or a truncated EnvΔ712 lacking the cytoplasmic tail. Cell lysates and virus particles were prepared and Western blotted as described above. The truncated form of gp41 in the Δ712 Env is indicated with an arrow. (D) 293T cells were transfected with the pNL4-3.AD8 IMC that is unmodified or modified with stop codons in the genes encoding reverse transcriptase (RT), RNAse H, integrase (IN), Vif, Vpr, Vpu or Nef. The cell lysates and virus particles were prepared and Western blotted as described above. Equal volumes of the clarified cell supernatants were used to infect TZM-bl cells for 48 hours, after which cells were lysed and the luciferase activity was measured. The results are representative of those obtained in two independent experiments, with the means and standard deviations reported. The full-length pNL4-3.AD8 IMC used in A-C encodes an AD8 Env with Bam (S752F I756F) changes in the cytoplasmic tail (see Fig. 3 below).

Next, we examined the effects of Env cleavage and cytoplasmic tail truncation on the quantity and quality of Env on virions produced by an IMC. Although the cleavage-defective AD8 Env was expressed well in the IMC-transfected cells, the uncleaved Env was incorporated into virions less efficiently than the cleavage-competent AD8 Env (Fig. 1C). Complete truncation of the Env cytoplasmic tail at residue 712 did not affect the gp120:gp160 ratio on virions, but slightly increased the level of cleaved Env incorporated.

To evaluate whether HIV-1 proteins that are not required for virion production affect Env incorporation, we individually knocked out the expression of reverse transcriptase (RT), RNAse H, integrase (IN), Vif, Vpr, Vpu and Nef (Fig. 1D). None of these knockouts affected the level of Env incorporation into VLPs or Env cleavage on the VLPs. As expected (140–142), the infectivity of the virus particles was dramatically reduced by the introduction of stop codons in the RT, RNAse H and IN open reading frames.

### Characterization of the AD8 Env on virus particles

Having established the suitability of the IMC expression system for producing VLPs, we characterized the AD8 Env on the virus particles. Deglycosylation with PNGase F and Endoglycosidase Hf demonstrated that all the cleaved Env on virus particles is modified by complex glycans; some of the gp160 glycoprotein on virus particles is also modified by complex carbohydrates (Fig. 2A). Therefore, all the cleaved Env and some of the uncleaved Env on these virus particle preparations has passed through the Golgi network (7). Interestingly, we observed two distinct bands for the deglycosylated gp160 and gp41 proteins, indicating the presence of truncated forms of these Envs. In parallel studies, we determined that clipping of the Env cytoplasmic tail at Arg 747 by the viral protease generates these truncated gp160 and gp41 proteins in the virus particles, a phenomenon that has been previously reported (143, 144). This protease-mediated clipping is enhanced by changes introduced into the AD8 Env cytoplasmic tail (F752S and F756I) as a result of the chimeric AD8-NL4-3 Env junction (at the BamHI site of *env*). Reverting these two cytoplasmic tail changes to the phenylalanine residues at 752 and 756 found in the wild-type HIV-1_AD8_ Env minimizes cytoplasmic tail clipping in the virus particles (see below). The AD8 Env containing Phe 752 and Phe 756, herein designated AD8 Bam, serves as the “wild-type” Env in this study.

**FIG 2.**
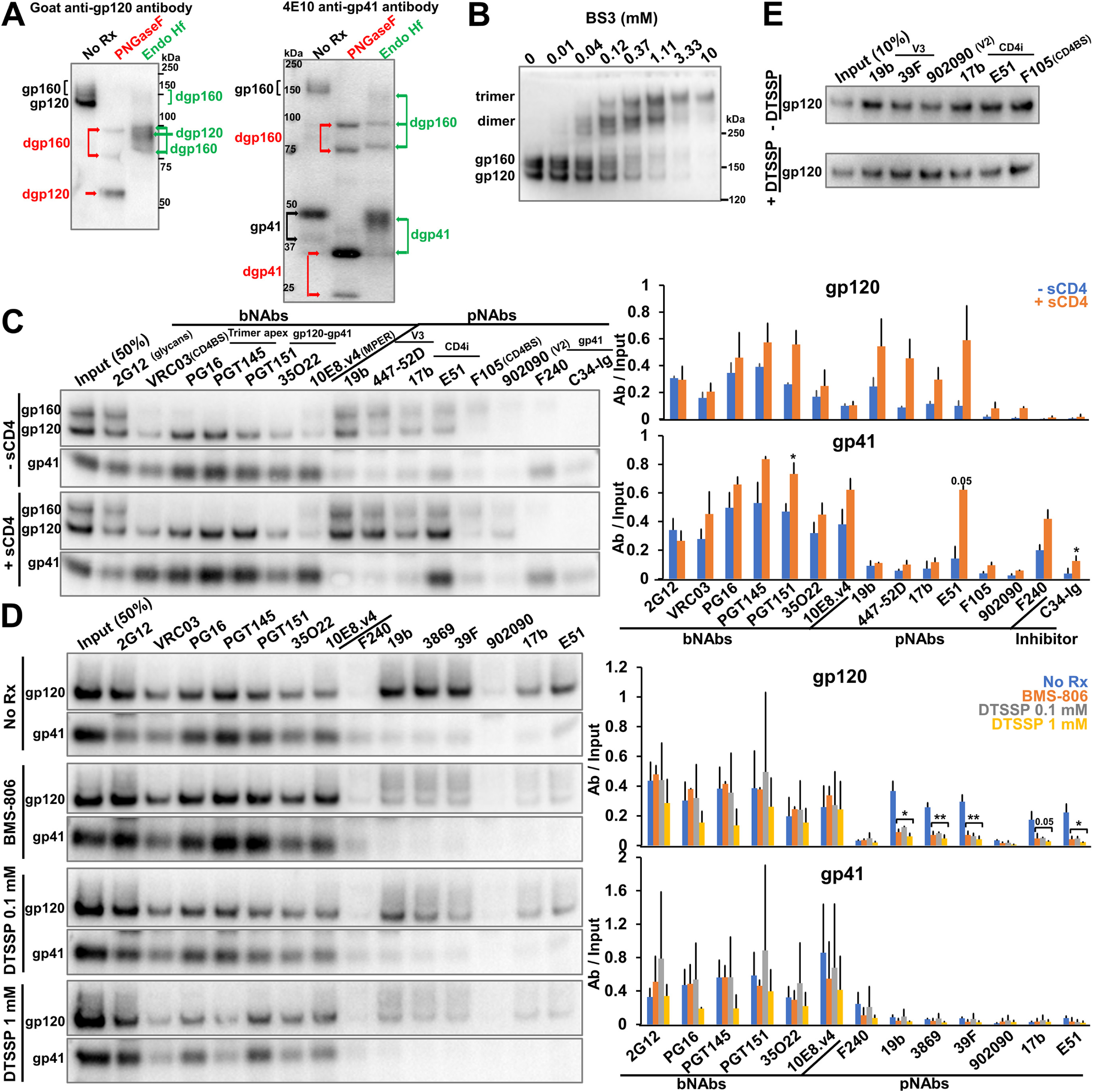
Characterization of the full-length AD8 Bam Env on IMC-produced virions. (A) 293T cells were transfected with the pNL4-3.AD8 IMC encoding an AD8 Env with Bam (S752F I756F) changes in the cytoplasmic tail (see Fig. 3 below). Forty-eight to seventy-two hours later, the cell supernatants were collected, filtered through a 0.45-μm membrane and centrifuged at 14,000-100,000 x g for 1 h at 4°C. Virus pellets were lysed, denatured and treated with PNGase F or Endo Hf for 1.5 h at 37°C, and Western blotted with a goat anti-gp120 antibody and the 4E10 anti-gp41 antibody. The deglycosylated (dg) Envs produced by PNGase F and Endo Hf are indicated by red and green labels, respectively. (B) Purified virus particles with AD8 Bam Envs were incubated with the BS3 crosslinker at the indicated concentrations for 30 min at room temperature. The samples were subsequently quenched, analyzed by reducing SDS-PAGE and Western blotted with a goat anti-gp120 antibody. (C) Purified virus particles were incubated with a panel of broadly neutralizing antibodies (bNAbs), poorly neutralizing antibodies (pNAbs) and the anti-HR1 C34-Ig peptide for 1 h at room temperature in the presence or absence of 10 μg/mL four-domain soluble CD4 (sCD4). The virus-antibody mixture was diluted twenty-fold with 1X PBS and centrifuged. The virus-antibody pellet was lysed and precipitated with Protein A-agarose beads for 1 h at 4°C. The beads were washed three times and Western blotted with a goat anti-gp120 antibody and the 4E10 anti-gp41 antibody. (D) Purified virus particles were incubated with 10 μM BMS-806 or with the indicated concentration of DTSSP crosslinker for 30 min at room temperature before the reactions were quenched with 100 mM Tris-HCl, pH 8.0. Env antigenicity on these virus particles was studied as described in (C). (E) 293T cells were transfected with the pNL4-3.AD8 Bam IMC expressing a soluble version of gp120. Forty-eight hours later, 0.45-μm filtered supernatant containing the soluble gp120 was crosslinked with 1 mM DTSSP as described above. Aliquots were then incubated with a panel of pNAbs and Protein A-agarose beads for 2 h at room temperature before the beads were washed and Western blotted with a goat anti-gp120 antibody. The results shown are representative of those obtained in two independent experiments. The means and standard deviations of the results in C and D are reported in the bar graphs in the panels on the right. The significance of the difference in antibody binding between treated and untreated samples was evaluated by a Student’s t test; *, p < 0.05; **, p < 0.01.

To examine the oligomeric composition of the AD8 Bam Envs on virions, purified virus particles were incubated with the crosslinker bis(sulfosuccinimidyl)suberate (BS3). As the BS3 concentration was increased, the AD8 Bam Env was efficiently crosslinked into gel-stable trimers, with no higher-molecular-weight forms observed (Fig. 2B). We conclude that essentially all the AD8 Bam Envs on these virus particles are assembled into trimers.

To examine the antigenicity of the AD8 Bam Env on virus particles, an immunoprecipitation assay was performed with a panel of broadly neutralizing antibodies (bNAbs) and poorly neutralizing antibodies (pNAbs) targeting the gp120 and gp41 glycoproteins. The bNAbs included 2G12 against gp120 outer-domain glycans (145), VRC03 against the gp120 CD4-binding site (CD4BS) (146), PG16 and PGT145 against quaternary gp120 epitopes at the trimer apex (147, 148), PGT151 and 35O22 against the gp120-gp41 interface (149, 150), and 10E8.v4 against the gp41 membrane-proximal external region (MPER) (106). The pNAbs included 19b and 447-52D, 3074, 3869 and 39F against the gp120 V3 region (151, 152), 17b and E51 against gp120 CD4-induced (CD4i) epitopes (153, 154), F105 against the gp120 CD4BS (155, 156), 902090 against the gp120 V2 region (157), and F240 against a Cluster I epitope on gp41 (158). C34-Ig is a fusion protein consisting of an immunoglobulin heavy chain and a gp41 HR2 peptide, C34, that targets the gp41 HR1 coiled coil (21). In the antigenicity assay performed in the absence of the soluble CD4 (sCD4) receptor, 2G12 precipitated both uncleaved and cleaved Envs, whereas the other bNAbs preferentially recognized the cleaved Envs (Fig. 2C, −sCD4 panels). All the bNAbs efficiently precipitated both gp120 and gp41 subunits, indicating that the subunits of the AD8 Bam Env trimers recognized by bNAbs remain associated during detergent solubilization, immunoprecipitation and washing of the immunoprecipitates. The pNAbs recognized the uncleaved gp160 Env, as expected (38, 39). The pNAbs against the gp120 V3 region and CD4i epitopes also precipitated the gp120 subunit of the cleaved AD8 Bam Env with surprising efficiency. Of interest, these pNAbs (19b, 447-52D and 17b) coprecipitated gp41 less efficiently than the gp120-directed bNAbs. The F240 antibody and C34-Ig recognized gp41 but did not coprecipitate gp120; these ligands are presumably recognizing gp41 glycoproteins that are not stably associated with gp120.

Together these observations suggest that the unliganded AD8 Bam Env on virus particles was processed in the Golgi apparatus and forms trimers with the following antigenic properties: (i) the uncleaved Env could be recognized by pNAbs, consistent with its conformational flexibility (39–45); (ii) the cleaved Envs could be recognized by bNAbs and some V3- and CD4i-directed pNAbs, which recognize gp120 structures involved in coreceptor binding (151-154,159,160); and (iii) the cleaved Envs that are recognized by bNAbs exhibit stronger gp120-trimer association in detergent than those that are recognized by pNAbs, suggesting differences in conformation between the two cleaved Env populations.

To examine the effects of receptor binding on the conformation of the AD8 Bam Env on virus particles, we performed the antigenicity assay in the presence of soluble four-domain CD4. In these experiments, we utilized conditions that were predetermined to allow detection of Env conformational changes with only minimal shedding of gp120 (data not shown). Incubation with sCD4 led to a significant increase in the binding of V3- and CD4i-directed pNAb binding to gp120; this was not accompanied by an increase in gp41 precipitation for the 19b, 447-52D and 17b pNAbs (Fig. 2C, +sCD4 panels). The E51 CD4i pNAb precipitated more gp120 and gp41 in the presence of sCD4, suggesting that complexes of the AD8 Bam Env, sCD4 and E51 antibody remain associated throughout detergent solubilization and washing. Mild increases in gp41 recognition by F240 and C34-Ig were also observed following incubation with sCD4, consistent with sCD4-induced shedding of gp120 from the Env trimers (161).

The small-molecule HIV-1 entry inhibitor, BMS-806, has been suggested to stabilize a pretriggered (State-1) conformation in membrane Envs (12,21,38,95,118,162,163). Chemical crosslinkers like 3,3’-dithiobis(sulfosuccinimidyl)propionate (DTSSP) are often used to limit the conformational dynamics of proteins (39). To examine the effects of these treatments on the conformation of the AD8 Bam Env on virus particles, we performed the immunoprecipitation assay in the presence of 10 µM BMS-806 or after crosslinking the viruses with 0.1 mM or 1 mM DTSSP (Fig. 2D). There were two significant differences in the antigenicity of Env on treated and untreated virus articles. First, both BMS-806 treatment or DTSSP crosslinking significantly reduced gp120 recognition by pNAbs. Crosslinking with 1 mM DTSSP did not affect pNAb binding to soluble gp120 monomers (Fig. 2E), ruling out an effect of DTSSP on the pNAb epitopes *per se* and indicating the importance of the Env trimer context to the observed reduction in pNAb binding. The results are consistent with BMS-806 and DTSSP treatment reducing spontaneous Env transitions from a pretriggered (State-1) conformation to more open downstream conformations recognizable by V3 and CD4i pNAbs. Second, treatment with the higher concentration of DTSSP led to a reduction in the binding of the PG16 and PGT145 bNAbs, which recognize V2 quaternary structures at the trimer apex (92,147,148,164). The binding of these bNAbs was not decreased by treatment with BMS-806 or lower concentrations of DTSSP. The observed reduction in PG16 and PGT145 bNAb binding after treatment with higher DTSSP concentrations may result from modification of key gp120 lysine residues shown to be important for the binding of these antibodies (92, 164).

### Effects of cytoplasmic tail clipping on Env conformation

As mentioned above, clipping of the AD8 Env cytoplasmic tail by the HIV-1 protease is enhanced by the alteration of two phenylalanine residues (Phe 752 and Phe 756) near the cleavage site (our unpublished observations). To evaluate the effect of cytoplasmic tail clipping on Env conformation, we compared the AD8 Env with the AD8 Bam Env, in which the S752F + I756F reversions were introduced. Little gp160 or gp41 clipping was detected in the lysates of cells expressing the AD8 and AD8 Bam Envs (Fig. 3A), consistent with this clipping being mediated by the HIV-1 protease, which is activated by dimerization in virion particles (165). On virus particles, approximately 72% of the AD8 gp41 was clipped, whereas only 28% of the AD8 Bam gp41 was clipped (Fig. 3A). Despite these differences in the level of clipped gp41, no significant differences were observed in the sensitivity of the AD8 and AD8 Bam viruses to neutralization by bNAbs, pNAbs, sCD4-Ig or BNM-III-170, a CD4-mimetic compound (CD4mc) (166) (Fig. 3B). The qualitative pattern of bNAb and pNAb binding to the AD8 and AD8 Bam virus particles was similar (Fig. 3C). The AD8 Env conformation and neutralization sensitivity are not apparently affected by protease-mediated gp41 clipping in virions.

**FIG 3.**
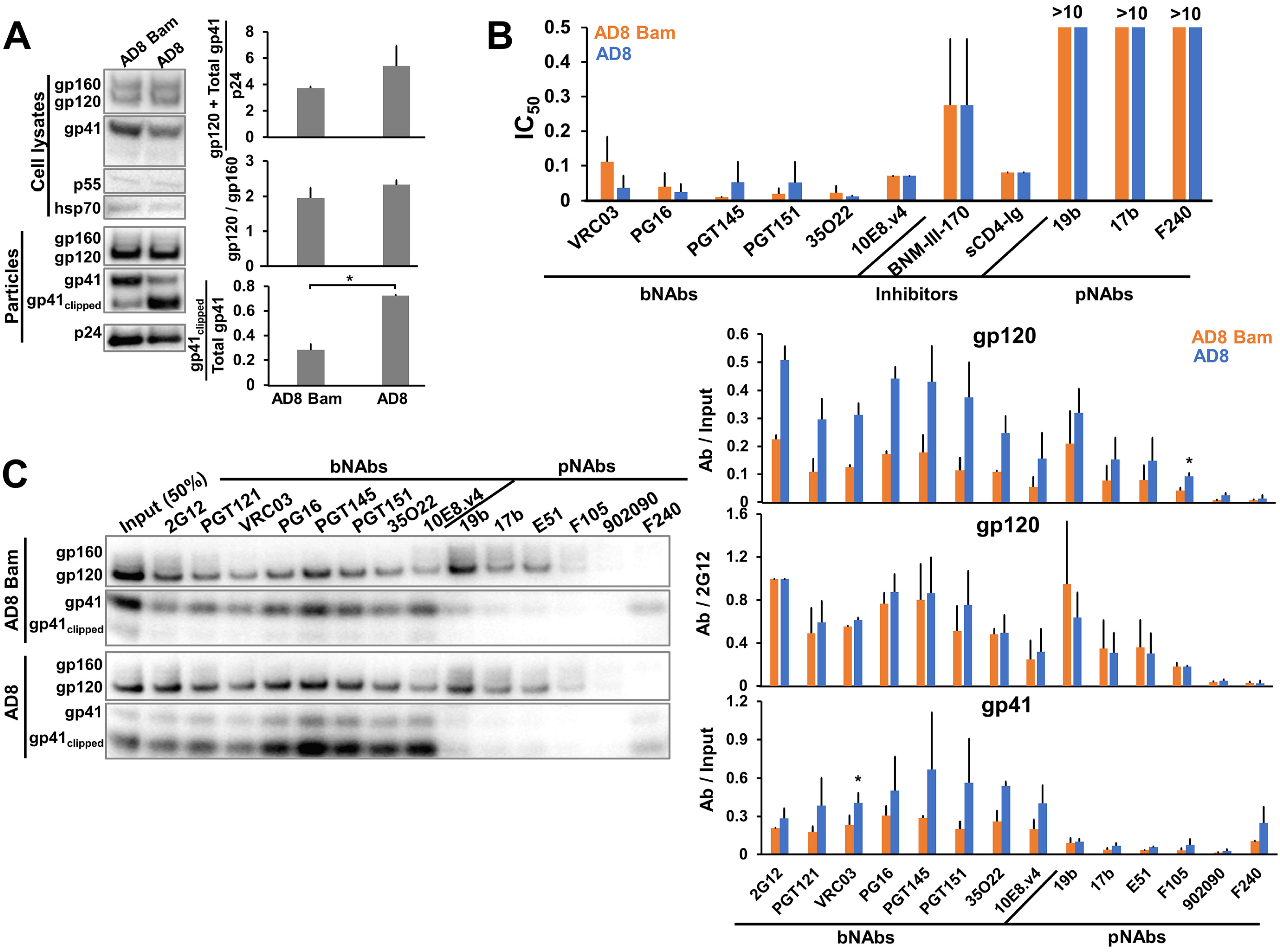
Effects of cytoplasmic tail clipping on Env conformation. (A) 293T cells were transfected with the pNL4-3.AD8 or pNL4-3.AD8 Bam IMCs, the latter encoding the AD8 Env with Bam (S752F I756F) changes in the cytoplasmic tail. Forty-eight to seventy-two hours later, the cell lysates and virus particles were prepared and Western blotted as described in the Fig. 1A legend. Clipping of the gp41 subunit in the virus particles was reduced by the Bam changes. (B) Seventy-two hours after transfection of 293T cells with IMCs, the cell supernatants were collected, clarified with a soft spin and incubated for 1 h at 37°C with a panel of bNAbs and pNAbs, soluble CD4-Ig or the CD4-mimetic compound BNM-III-170. The mixture was added to TZM-bl cells for 48 h, after which cells were lysed and the luciferase activity measured. The 50% inhibitory concentrations of the Env ligands are reported in μg/mL except for BNM-III-170 (in μM). (C) Antigenicity of the VLP AD8 Env with and without the Bam changes was analyzed as described in the Fig. 2C legend. The results are representative of those obtained in two independent experiments. The means and standard deviations of the results in B and C are reported in the bar graphs. The significance of the difference in antibody binding to the AD8 and AD8 Bam Envs in C was evaluated by a Student’s t test; *, p < 0.05.

### Effects of State-1-stabilizing and -destabilizing changes on virion Env

Previous studies have identified HIV-1 Env variants in which the pretriggered (State-1) conformation is stabilized, rendering the viruses more resistant to cold inactivation and to inhibition by sCD4 and the CD4mc BNM-III-170 (167). Conversely, Env variants in which State 1 is destabilized often exhibit global increases in sensitivity to inactivation at 0° C and to neutralization by sCD4, CD4mcs and pNAbs (107-113,168-169). To evaluate the effect of State-1-stabilizing and -destabilizing changes on the antigenicity of Env on viruses, we introduced these changes into the AD8 Bam Env. Three changes (Q114E, Q576K and A582T) critical to the State-1-stabilized phenotype of a previously reported HIV-1_AD8_ Env variant, AE.2 (167), were introduced into the AD8 Bam Env to create the Tri Bam Env. Multiple Env polymorphisms found in the AE.2 Env, including the State-1-stabilizing A114E, Q567K and A582T changes, were introduced into the AD8 Bam Env to create the AE.1 Bam Env (see Materials and Methods for details). The sensitivity of viruses with the AD8 Bam, Tri Bam and AE.1 Bam Envs to inhibition by sCD4-Ig and the CD4mc BNM-III-170 was measured using TZM-bl target cells. The sCD4-Ig IC_50_ values were 0.076 (AD8 Bam), >20 (Tri Bam) and >20 (AE.1 Bam) µg/ml. The BNM-III-170 IC_50_ values were 0.22 (AD8 Bam), >100 (Tri Bam) and >100 (AE.1 Bam) µM. The viruses with the Tri Bam and AE.1 Bam Envs were also more resistant to inactivation on ice than viruses with the AD8 Bam Env; after 1 day on ice, viruses with the AD8 Bam Env lost ∼45% of their infectivity, whereas the infectivity of the viruses with Tri Bam and AE.1 Bam Envs was unaffected (data not shown). These phenotypes are consistent with State-1 stabilization of the Tri Bam and AE.1 Bam Envs, relative to the AD8 Bam Env (48,113,167).

State-1-destabilizing changes (N197S in gp120, NM 625/626 HT and D674N in gp41) (107) were introduced into the AD8 Bam Env to create the AD8 Bam 197 HT N Env. The AD8 Bam 197 HT N virus was neutralized by the 19b and 17b pNAbs, whereas the AD8 Bam virus was not (Fig. 4A). Both viruses were neutralized comparably by bNAbs. Relative to the AD8 Bam virus, the AD8 Bam 197 HT N virus was inhibited more effectively by BNM-III-170. These phenotypes are consistent with State-1 destabilization of the AD8 Bam 197 HT N Env relative to the AD8 Bam Env.

**FIG 4.**
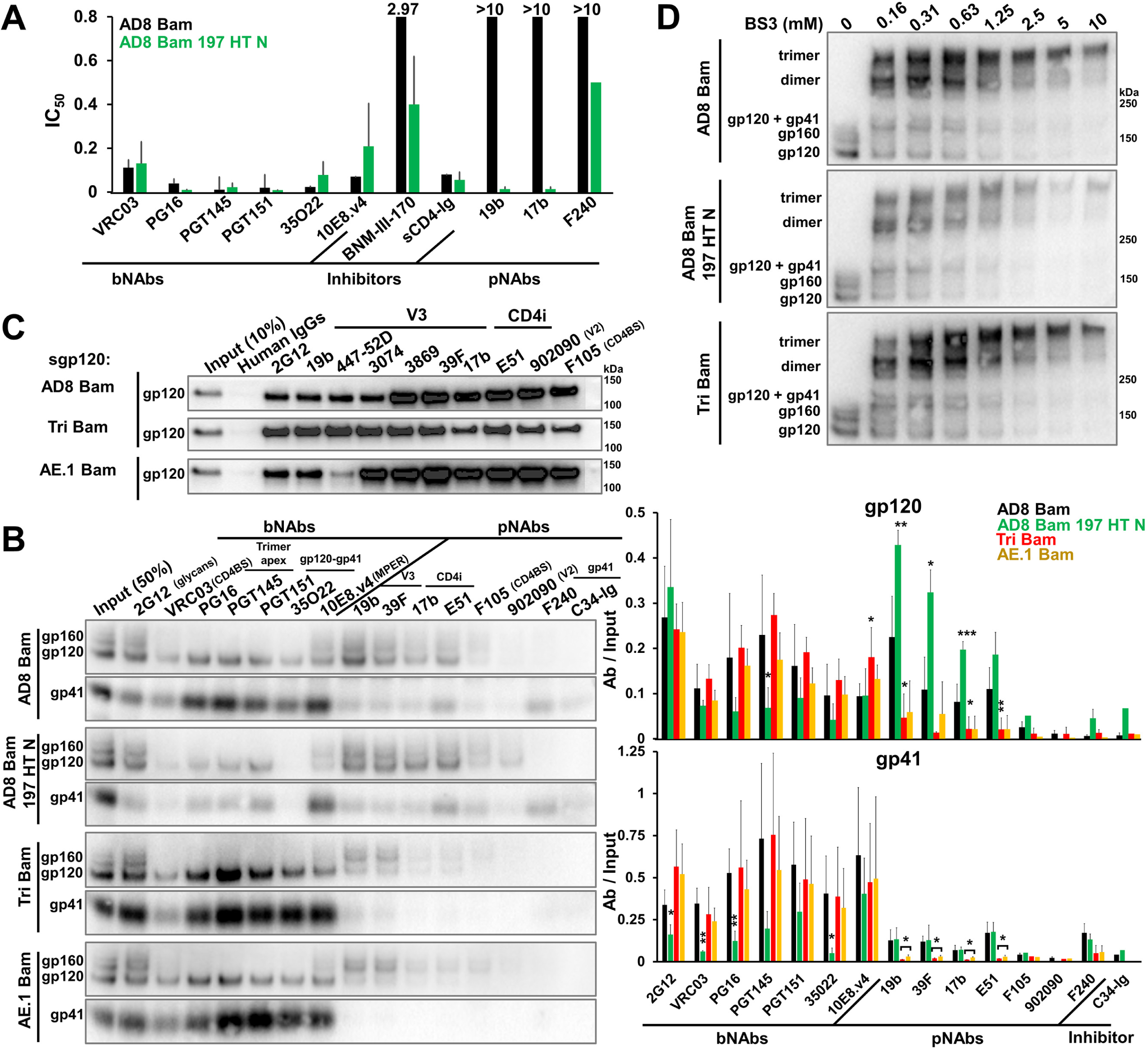
Effects of State-1-stabilizing and -destabilizing changes on virion Env. (A) Neutralization of the AD8 Bam and AD8 Bam 197 HT N viruses by the indicated Env ligands was measured as described in the Fig. 3B legend. The 50% inhibitory concentrations of the Env ligands are reported in μg/mL except for BNM-III-170 (in μM). (B) 293T cells were transfected with pNL4-3.env IMCs expressing the AD8 Bam Env, the State-1-destabilized AD8 Bam 197 HT N Env and the State-1-stabilized Envs Tri Bam and AE.1 Bam. Seventy-two hours later, virions were purified and the antigenicity of Env on virus particles was analyzed as described in the Fig. 2C legend. (C) 293T cells were transfected with the pNL4-3.env IMCs expressing soluble versions of gp120 (sgp120) from the indicated Envs. Forty-eight hours later, the cell supernatant containing secreted gp120 was collected, filtered through a 0.45 μm membrane and incubated with the indicated antibodies and Protein A-agarose beads for 2 h at room temperature. The beads were washed and Western blotted with a goat anti-gp120 antibody. (D) Purified virus particles were incubated with the crosslinker BS3 at the indicated concentrations for 30 minutes at room temperature, after which the reactions were quenched and samples were analyzed by reducing SDS-PAGE and Western blotted with a goat anti-gp120 antibody. The results are representative of those obtained in two independent experiments. The means and standard deviations of the results in A and B are shown in the bar graphs. The significance of the difference in antibody binding between Env mutants and AD8 Bam Env was evaluated by a Student’s t test; *, p < 0.05; **, p < 0.01; ***, p < 0.001.

Compared to the parental AD8 Bam Env, the State-1-stabilized Tri Bam and AE.1 Bam Envs on virus particles were recognized as well by bNAbs and, importantly, only weakly by the V3 and CD4i pNAbs (Fig. 4B). Soluble forms of the gp120 glycoproteins from the AD8 Bam, Tri Bam and AE.1 Bam Envs were precipitated by these pNAbs efficiently (Fig. 4C), indicating that the observed reduction in pNAb binding to the State-1-stabilized Envs on virus particles was not a result of disruption of the epitopes. As a control in this experiment, the recognition of the AE.1 gp120, which contains an R315K change in the V3 region (167), by the 447-52D pNAb was decreased. The R315K change does not affect the binding of the other V3-directed pNAbs (47, 170) used in our study.

Interestingly, the antigenicity of the AD8 Bam 197 HT N Env displayed the opposite trends. Compared with the parental AD8 Bam Env, the AD8 Bam 197 HT N Env was recognized less well by bNAbs (PG16, PGT145 and 35O22) with epitopes dependent on quaternary Env structure (Fig. 4B). Conversely, recognition of the AD8 Bam 197 HT N Env by V3 and CD4i pNAbs was relatively increased.

To examine whether the antigenic differences between these State-1-stabilized and -destabilized Envs might be influenced by the oligomeric state of the Env variants, virus particles containing the Env variants were crosslinked with the BS3 crosslinker. The parental AD8 Bam, State-1-destabilized AD8 Bam 197 HT N and State-1-stabilized Tri Bam envelope glycoproteins were crosslinked to trimers with equal efficacy (Fig. 4D).

To summarize, HIV-1 Envs with varying degrees of State-1 stability retain trimeric configurations on virus particles. The spontaneous exposure of V3 and CD4i pNAb epitopes on the cleaved Env trimer is inversely related to the degree of State-1 stability. State-1-stabilized Envs effectively maintain the epitopes recognized by bNAbs, particularly those dependent on quaternary conformation. By contrast, the State-1-destabilized Env was recognized inefficiently by bNAbs.

### Shedding of gp120 from virus particles

The non-covalent association of gp120 with the Env trimer is prone to disruption, either spontaneously or in response to the binding of ligands (sCD4, CD4-mimetic compounds) that induce Env conformations downstream of State 1 (137-139,161,171). We set out to study the shedding of gp120 from Envs with different degrees of State-1 stability on virus particles produced by IMCs. To establish optimal assay conditions, we measured gp120 shedding from the AD8 Bam Env after a one-hour incubation with different concentrations of sCD4 or the CD4mc BNM-III-170 at various temperatures (Fig. 5A). Both sCD4 and BNM-III-170 induced gp120 shedding in a dose-dependent manner. For both sCD4 and BNM-III-170, shedding of gp120 was slightly more efficient at lower temperatures than at 25° or 37°C. At the highest sCD4 and BNM-III-170 concentrations tested, ∼6-10% of the total gp120 on the virus particles was shed after a 1-hour incubation at 0° or 4°C.

**FIG 5.**
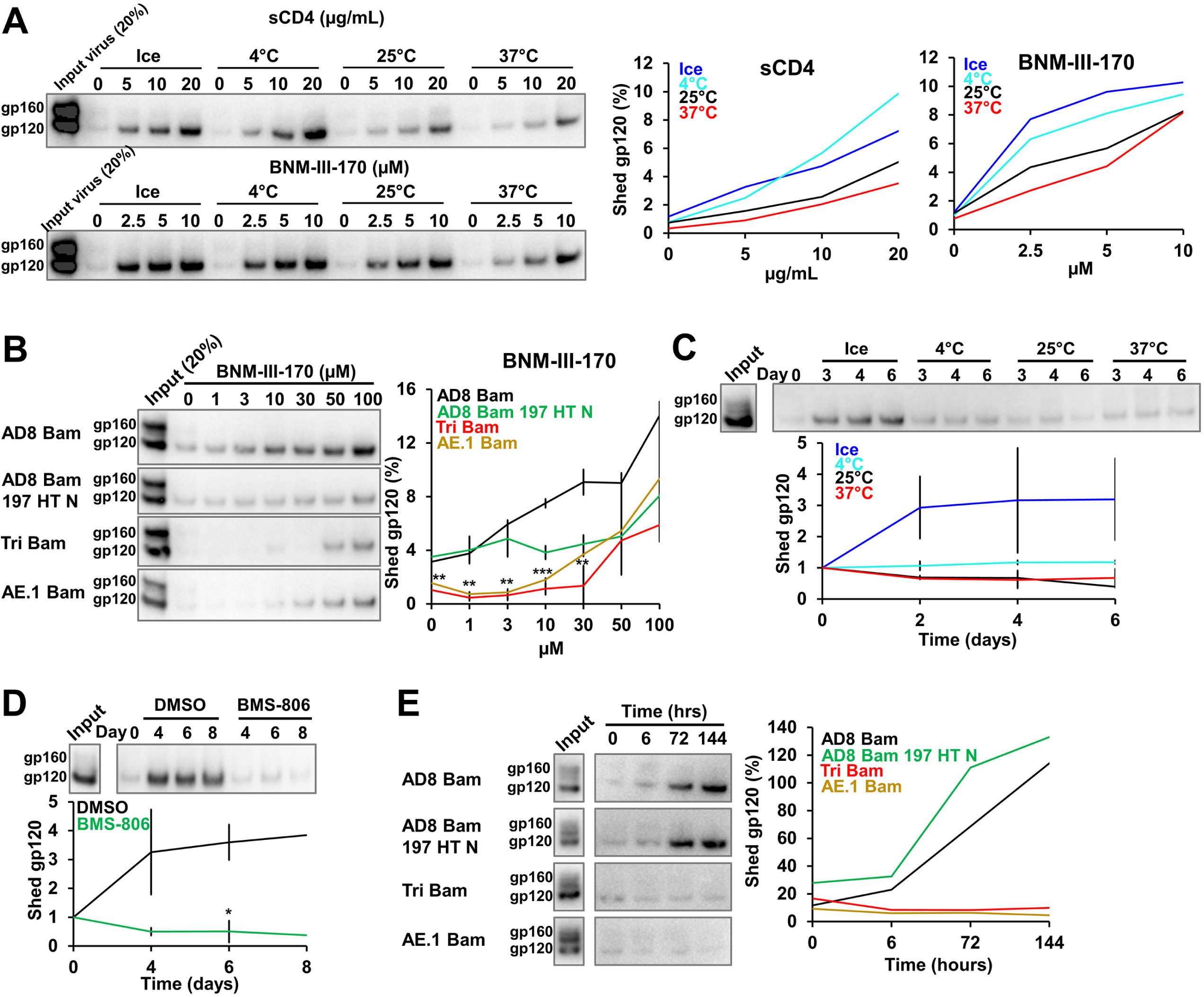
Shedding of gp120 from Env on virus particles. (A) 293T cells were transfected with the pNL4-3.AD8 Bam plasmid. Forty-eight to seventy-two hours later, the cell supernatants were collected, filtered through a 0.45-μm membrane and centrifuged at 100,000 x g for 1 h at 4°C. Virus pellets were resuspended and incubated with four-domain soluble CD4 (sCD4) or the CD4-mimetic compound BNM-III-170 at the indicated concentrations and temperatures for 1 h. Virus particles were again pelleted and the supernatants containing shed gp120 were incubated with GNL beads for 2 h at room temperature. Beads were washed and Western blotted with a goat anti-gp120 antibody. The percentage of the gp120 Env on the input virus that was detected in the supernatants is plotted in the graphs on the right. (B) Shedding of gp120 from different Envs after a 1-h room temperature incubation with the indicated concentrations of BNM-III-170 was analyzed as described in A. (C) Purified virus particles were incubated at different temperatures for different lengths of time. Shed gp120 was then analyzed as described in A. (D) Purified virus particles with the AD8 Bam Env were incubated on ice in the presence of DMSO or 10 μM BMS-806 for different lengths of time. Shed gp120 was then analyzed as described in A. (E) Purified virus particles containing different AD8 Env variants were incubated on ice for the indicated lengths of time and shed gp120 was then analyzed as described in A. Except for A, the results shown are representative of those obtained in two independent experiments. In B-D, the means and standard deviations are reported. The significance of the difference between DMSO- and BMS-806-treated samples (C) or between Tri Bam and AE.1 Bam Envs compared to AD8 Bam Env (D) was evaluated by a Student’s t test; *, p < 0.05; **, p < 0.01; ***, p < 0.001.

The resistance of closely matched HIV-1 Env variants to inhibition by CD4mcs is a valuable indicator of the degree of State-1 stabilization (167). To determine whether CD4mc-induced gp120 shedding correlates with this functional phenotype, we compared BNM-III-170-induced shedding of gp120 from virus particles with the AD8 Bam, AD8 Bam 197 HT N, Tri Bam and AE.1 Bam Envs (Fig. 5B). Both State-1-stabilized Envs, Tri Bam and AE.1 Bam, were less sensitive to gp120 shedding induced by BNM-III-170 than the AD8 Bam Env.

In HIV-1 variants with closely matched Envs, resistance to cold inactivation is another useful indicator of the degree of State-1 stabilization (167,172,173). We sought to evaluate the relationship between Env conformation and the spontaneous shedding of gp120 at different temperatures. In initial experiments, we incubated virus particles with the AD8 Bam Env at different temperatures for various times and measured the amount of shed gp120 (Fig. 5C). Shedding of gp120 was significantly greater after incubation of the viruses on ice than at higher temperatures. Thus, the inactivation of the infectivity of viruses with the AD8 Bam Env by incubation on ice coincides with destabilization of the Env trimer and loss of the gp120 subunit.

To evaluate whether HIV-1 Envs stabilized in a pretriggered (State-1) conformation could better resist cold-induced gp120 shedding, we took two approaches. First, we incubated viruses with the AD8 Bam Env on ice in the absence or presence of BMS-806, which stabilizes the State-1 Env conformation (12,21,38,95,118). Treatment with BMS-806 effectively prevented gp120 shedding from the AD8 Bam Env even after an 8-day incubation on ice (Fig. 5D). Second, we compared gp120 shedding at 0° C for viruses with State-1-stabilizing and -destabilizing changes in Env. The State-1-stabilized Envs, Tri Bam and AE.1 Bam, retained gp120 even after a 6-day incubation on ice, whereas the parental AD8 Bam and the State-1-destabilized AD8 Bam 197 HT N Envs shed much of their gp120 subunits during this time period (Fig. 5E).

To summarize, incubation at 0°C leads to destabilization of the HIV-1 AD8 Bam Env trimer on virus particles, resulting in gp120 shedding. Envs stabilized in a pretriggered (State-1) conformation are better able to resist the trimer-destabilizing effects of exposure to cold. Measurements of gp120 shedding from virus particles correlate well with virus functional phenotypes and can serve as useful indicators of Env conformational state, even for Envs with limited ability to support HIV-1 infection.

### Effects of 0**°**C incubation on the antigenicity of a State-1-stabilized Env

Although the State-1-stabilized Tri Bam Env did not shed gp120 after prolonged incubation on ice, other more subtle changes to the conformation of the Tri Bam Env trimer might have resulted from cold exposure. To address this, we compared the antigenicity of the Tri Bam Env after a 7-day incubation on ice with that of a Tri Bam Env not incubated on ice (Fig. 6). Exposure to 0°C for 7 days had little effect on the amount of Tri Bam Env on the virus particles or on its recognition by bNAbs or pNAbs. We note that the V3 and CD4i pNAbs mainly recognize the uncleaved Tri Bam Env on the virus particles, and any low-level recognition of gp120 by these antibodies was not accompanied by coprecipitation of gp41. We conclude that the effects of the Q114E, Q567K and A582T changes on Env conformation allow the virion Env trimers to withstand cold stress for at least one week.

**FIG 6.**
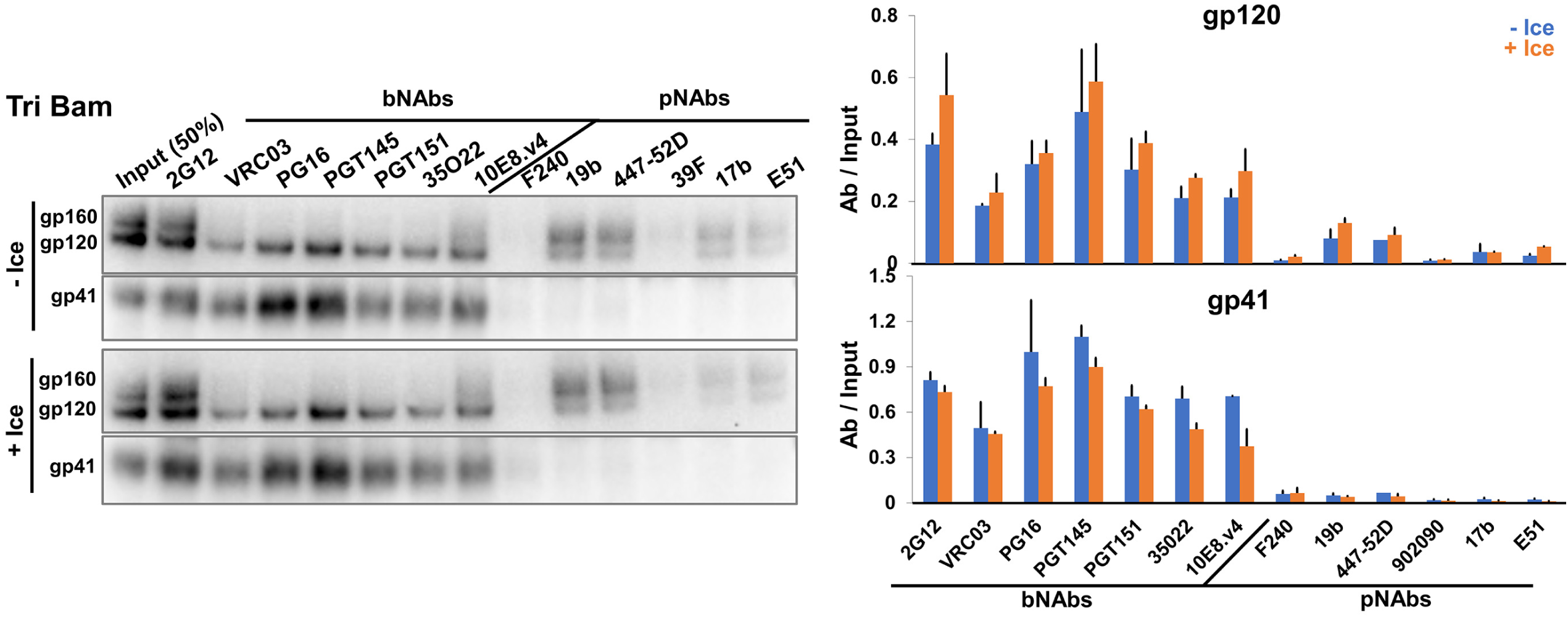
Maintenance of Tri Bam Env antigenicity on virions after prolonged incubation on ice. HEK 293T cells were transfected with the pNL4-3.Tri Bam infectious molecular clone. Seventy-two hours later, virus particles were purified, aliquoted and stored at −80°C. Pilot experiments found no detectable difference in the Env content or antigenicity of VLPs that were analyzed directly after production or were frozen at −80°C once. One −80°C aliquot was thawed and incubated on ice for 7 days (+Ice). At this time, a second aliquot was thawed to serve as a reference control (−Ice). Both samples were pelleted and washed to remove any shed gp120, after which the Env antigenicity on virus particles was analyzed as described in the Fig. 2C legend. The results are representative of those obtained in two independent experiments, with the means and standard deviations reported in the bar graphs on the right.

### Characterization of virion Envs from other HIV-1 strains

The studies above utilized closely matched HIV-1_AD8_ Env variants with amino acid residue changes that specifically alter Env triggerability (107, 167); analysis of these variants identified useful phenotypic indicators of the degree of State-1 stabilization. To examine whether these conformational indicators could be generalized to Envs from other HIV-1 strains, we studied the laboratory-adapted, tier-1 HIV-1_NL4-3_ (clade B); the primary, tier-2/3 HIV-1_JR-FL_ (clade B); and the primary, tier-2/3 HIV-1_BG505_ (clade A). HIV-1_JR-FL_ and HIV-1_BG505_ Envs are of particular interest because pseudoviruses with these Envs exhibit levels of resistance to cold inactivation and CD4mc inhibition that are comparable to those of pseudoviruses with the AD8 Env containing the State-1-stabilizing changes examined here (167). The JR-FL E168K Env variant was used in the analysis of Env antigenicity as the E168K change in the gp120 V2 region allows recognition by the V2 quaternary bNAbs, PG16 and PGT145, without detectable effects on other properties of the HIV-1_JR-FL_ Env (92,147,148,164). Comparing the antigenicity of the Env variants on virus particles (Fig. 7A), we noted trends similar to those observed for the AD8 Bam Env variants: (i) the cleaved Env in the more triggerable NL4-3 Env was not efficiently precipitated by the bNAbs (PG16, PGT145, PGT151 and 35O22) that recognize epitopes dependent on quaternary Env structure; nonetheless, the NL4-3 virus was efficiently neutralized by the PG16 and PGT145 antibodies (data not shown); (ii) the cleaved JR-FL E168K Env was precipitated by all the bNAbs but, with the exception of weak precipitation by the 19b and 447-52D V3 pNAbs, not by most pNAbs; and (iii) although the level of cleaved BG505 Env on the virus particles was very low, this small amount of cleaved Env was detected better by the bNAbs than by pNAbs. Deglycosylation of the precipitated BG505 Env proteins confirmed the efficient recognition of gp120 by all the bNAbs tested, with detectable recognition of gp120 by the 19b and 447-52D V3 pNAbs and by the 17b and E51 CD4i pNAbs (data not shown). We confirmed that the pNAb epitopes are present on the gp120 glycoproteins of these three HIV-1 strains (with the exception of the 19b V3 epitope on the NL4-3 Env) (Fig. 7B).

**FIG 7.**
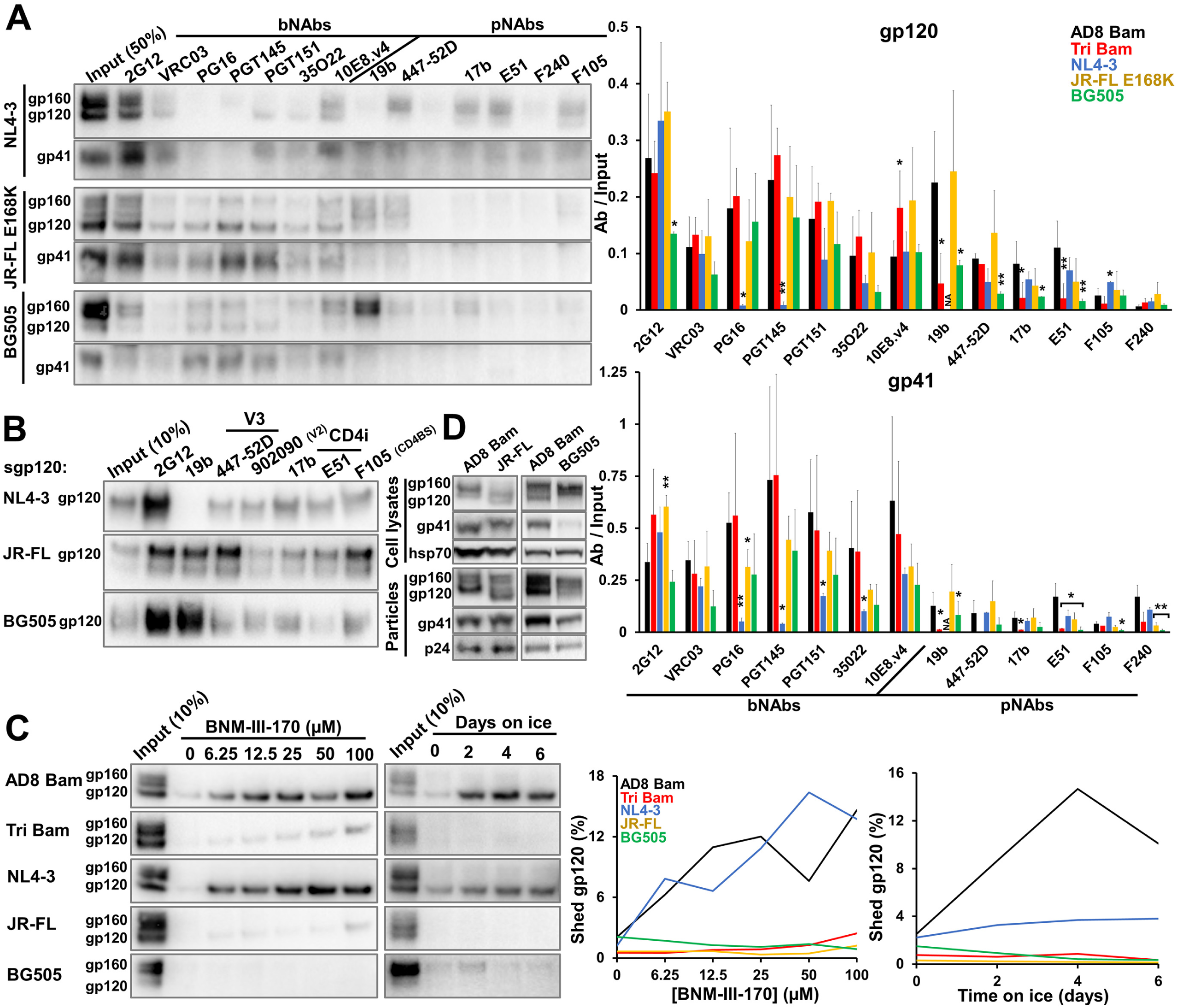
Characterization of VLP Envs from other HIV-1 strains. (A) 293T cells were transfected with pNL4-3.env infectious molecular clones expressing the NL4-3, JR-FL E168K and BG505 Envs. Seventy-two hours later, virus particles were purified and Env antigenicity on virus particles was analyzed as described in the Fig. 2C legend. Note that in A only, the JR-FL E168K mutant was used to allow recognition by V2 quaternary bNAbs (PG16, PGT145) (147, 148). The means and standard deviations of the results obtained in two independent experiments are reported in the bar graph at the right of the figure. The significance of the difference between AD8 Bam Env and other Envs was evaluated by a Student’s t test; *, p < 0.05; **, p < 0.01; NA, not applicable. (B) The antigenicity of soluble gp120 versions of the indicated Envs was performed as described in the Fig. 4C legend. (C) The susceptibility of VLP Envs to gp120 shedding induced by BNM-III-170 or ice incubation was evaluated as described in the Fig. 5B and 5E legend. (D) Env glycoprotein expression, processing and incorporation into virus particles were examined as described in the Fig. 1A legend. The results are representative of those obtained in at least two independent experiments.

We examined the shedding of gp120 from viral particles containing Envs from the different HIV-1 strains in response to the CD4mc BNM-III-170 and to ice exposure (Fig. 7C). The NL4-3 Env shed gp120 efficiently following incubation with BNM-III-170 or on ice. By contrast, the JR-FL and BG505 Envs shed gp120 minimally in response to the CD4mc or cold exposure. These gp120 shedding efficiencies correlate with the susceptibility of these viruses to cold inactivation or inhibition by BNM-III-170 (167). Thus, the antigenicity and gp120 shedding assays using IMC-generated virus particles revealed important distinctions among Envs with different triggerability levels, even for Envs derived from different HIV-1 strains and clades. Additionally, viruses produced from cells transfected with IMCs may demonstrate improved Env proteolytic processing in cases, e.g., the JR-FL and BG505 Envs, where low levels of Env cleavage have been observed in the transfected cells (Fig. 7D) (167). However, even when produced by an IMC, the BG505 Env still displayed a very low level of cleavage in cell lysates and on virus particles. Therefore, a high level of cleaved Env on virus particles is one criterion that could be used to prioritize HIV-1 strains for structural and immunogenicity studies of membrane Envs.

### Characterization of Env in virions produced from infected T cells

The above analyses were performed with virus particles produced transiently from transfected 293T cells. To examine whether the observed Env phenotypes would also be associated with virions produced from infected T cells, we analyzed the antigenicity of the AD8 Bam Env and the E.1 Bam Env in virions produced from infected C8166-R5 cells. C8166-R5 cells are human CD4^+^ T lymphocytes transformed by human T-cell leukemia virus (HTLV-I) and transduced with a vector expressing human CCR5 (174). The E.1 Bam Env is identical to the AE.1 Bam Env except that one of the State-1-stabilizing changes, A582T, has been reverted (167). The E.1 Bam Env was studied here instead of the Tri Bam or AE.1 Bam Envs because the E.1 Bam Env was more infectious in C8166-R5 cells, but nonetheless retained most State-1-stabilized phenotypes (reference 167 and data not shown). The AD8 Bam Env on infectious virions produced in C8166-R5 cells displayed an antigenic pattern similar to that of the virus particles produced from transfected 293T cells, i.e., gp120 was recognized by bNAbs and V3 and CD4i pNAbs, but the coprecipitation of gp41 was less efficient for the pNAbs than for the bNAbs (Fig. 8A). Relative to the antigenicity of AD8 Bam Env, the cleaved E.1 Bam Env on virions demonstrated strong bNAb binding and decreased pNAb binding. The consistency between the antigenicity of Envs on viruses produced from transfected 293T cells and an infected T cell line support the generality and intrinsic nature of the observed Env phenotypes.

**FIG 8.**
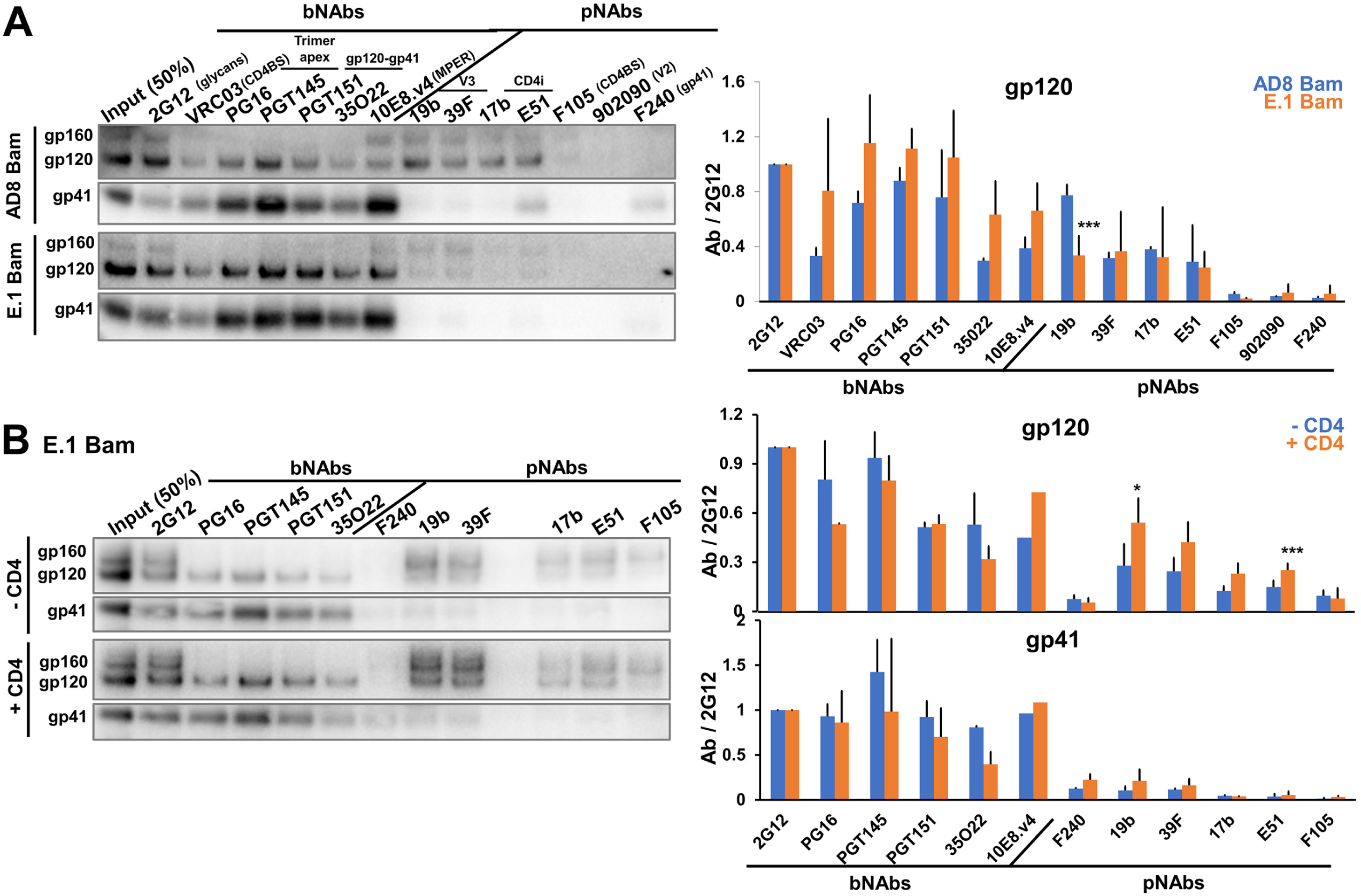
Characterization of Envs on virus particles from infected T cells. (A) HEK 293T cells were transfected with pNL4-3.env infectious molecular clones expressing the Strep-tagged AD8 Bam or E.1 Bam Envs. Addition of the Strep tag to the Env C terminus does not affect neutralization sensitivity nor Env antigenicity on virus particles (data not shown). Seventy-two hours later, the cell supernatants were clarified and ID_50_ values were determined using TZM-bl target cells. C8166-R5 cells were then infected with virus particles at a multiplicity of infection of 0.1. Cells were washed 5-16 h after infection and resuspended in fresh medium. Fresh medium was supplemented at three days after infection. Six to seven days after infection, virus particles were collected, purified and Env antigenicity was analyzed as described in the Fig. 2C legend. (B) 293T cells were transfected with the pNL4-3.E.1 Bam Strep infectious molecular clone with or without a CD4-expressing plasmid at a 1:0.1 weight ratio. Seventy-two hours later, virus particles were collected, purified and Env antigenicity was analyzed as described in the Fig. 2C legend. The results are representative of those obtained in at least two independent experiments. The means and standard deviations of the results from A and B are reported in the bar graphs (right panels). The significance of the difference between the wild-type AD8 Bam Env and E.1 Bam Env (A) or between VLP Envs made in the absence or presence of CD4 (B) was evaluated by a Student’s t test; *, p < 0.05; ***, p < 0.001.

In multiple repeat experiments, we noted that the binding of the cleaved E.1 Bam Env by V3 and CD4i pNAbs was variable and, in some cases, at a level comparable to that of the AD8 Bam Env (see average values in Fig. 8A, right panel). Because C8166-R5 cells express CD4 that could potentially interact with HIV-1 Env (175, 176), we tested whether CD4 coexpression could affect the conformation of a State-1-stabilized Env on virus particles. To that end, we transfected 293T cells with the E.1 Bam IMC with or without a plasmid expressing human CD4. Analysis of the E.1 Bam Env antigenicity on virus particles showed that coexpression of CD4 resulted in modest increases in the binding of V3 and CD4i pNAbs to gp120, without accompanying increases in the coprecipitation of gp41 (Fig. 8B). We conclude that coexpression of CD4 in cells producing virus particles can in some circumstances lead to an increased sampling of non-State-1 Env conformations on the viral particles.

## DISCUSSION

The development of an effective AIDS vaccine has been frustrated by the inefficiency with which current Env immunogens, including stabilized soluble trimers, elicit neutralizing antibodies with breadth against primary HIV-1 strains (64–75). The evolution of bNAbs during natural HIV-1 infection likely is driven by the mature (cleaved) State-1 Env trimer on viral or cell membranes. The association of Env with the membrane is important for maintaining a State-1 Env conformation and for the correct composition of Env glycans (11,90–93,102–113), both of which can potentially influence the binding of bNAbs and their precursors.

Virus-like particles (VLPs) offer a means to access native, functional HIV-1 Envs in a natural membrane context. Indeed, several groups have explored VLP Env composition, antigenicity, structure, dynamics and immunogenicity (12,95,118,121-137,177–179). Structural and conformational heterogeneity in VLP Envs complicates efforts to characterize the virion spike and to develop these membrane Envs as immunogens presenting a pretriggered (State-1) conformation to the immune system. One important source of conformational heterogeneity is the uncleaved gp160 Env, which is flexible and binds multiple pNAbs (40–45). In this study, we utilized infectious molecular clones (IMCs) to produce virus particles with sufficient levels of cleaved HIV-1 Env for detailed analysis of Envs on virus particles. In this respect, virus particles produced by IMCs were superior to pseudotyped viruses, where Env cleavage was inefficient and not sufficiently increased by lowering the amount of Env-expressing plasmid transfected with the Gag-expressing plasmid. Our results are consistent with those of previous studies (136, 180). In one such study (136), extremely low ratios (e.g., 1:80) of Env:Gag expressor plasmids were required to achieve a level of Env cleavage comparable to that produced by an IMC. At such low Env:Gag ratios, the low levels of Env on the VLPs create additional impediments to the characterization of the virion Envs. In addition to the use of IMCs, we also utilized Western blotting that could distinguish the phenotypes of cleaved and uncleaved Envs; this distinction is essential for characterizing the conformations of the different Env populations present on virus particles.

In contrast to pseudotyped viruses, IMC-produced virus particles generally exhibited levels of cleaved Env similar to those in virions produced from infected T cells. The infectivity of IMC-produced viruses can be inactivated by the introducing conservative changes that eliminate reverse transcriptase, RNAse H or integrase expression without compromising the enrichment of cleaved Env on the virus particles. Disruption of *vif*, *vpr*, *vpu* and *nef* also did not affect Env amount or level of cleavage on the virus particles (Fig.1D). Combinations of these inactivating mutations could be introduced into IMCs to minimize the possibility of infectious virus in VLP preparations. Vpu and Nef contribute to the down-regulation of CD4-Env complexes on the surface of infected cells and virions, and therefore could be of value in VLP-producing cells that express CD4 (176,181–185). However, we found that even with intact *vpu* and *nef* genes on the IMC, CD4 expression in the producer cell increased the exposure of CD4-induced pNAb epitopes on the VLP Env (Fig. 8B). Using CD4-negative producer cells avoids this potential problem. The HIV-1 protease, which was left intact on the IMCs to allow proteolytic maturation of the VLPs, can also clip the cytoplasmic tail of Env from some HIV-1 strains (143, 144). We found that this cytoplasmic tail clipping was enhanced by chimerism of the HIV-1_AD8_ Env construct near the cleavage site and could be remedied by changes near the junction sequences. Nonetheless, cytoplasmic tail clipping exerted no detectable effect on AD8 Env antigenicity or neutralization sensitivity. These results are consistent with previous studies showing that complete truncation of the HIV-1_AD8_ Env cytoplasmic tail does not detectably affect virus neutralization sensitivity (94).

IMC-produced virus particles with high levels of cleaved Env allowed direct analysis of the virion Env population, essentially all of which is trimeric. The antigenicity and gp120 shedding analyses revealed the existence of at least three populations of cleaved AD8 Env trimers (Fig. 9A):

### 1) Pretriggered (State-1) Env

This cleaved Env population is marked by its recognition by bNAbs but not pNAbs. This Env population is maintained after treatment with BMS-806 and crosslinkers and is increased by Env changes that stabilize the pretriggered conformation (167). Notably, as seen in the ability of gp120- or gp41-directed bNAbs to coprecipitate the other subunit, the Env trimers in this population are stable in detergent lysates. This Env population also is more resistant to gp120 shedding induced by incubation on ice or with CD4mc. Lower Env triggerability and increased State-1 occupancy are often associated with increased intersubunit interactions that stabilize the trimer (38, 167). In agreement with this, BMS-806-treated and State-1-stabilized Envs are relatively resistant to gp120 shedding after exposure to 0° C or detergent. The cleaved, State-1-stabilized Tri Bam Env on the virus surface maintained its antigenicity for at least one week on ice.

### 2) Envs in more open conformations

This cleaved Env population is marked by its recognition by V3 and CD4i pNAbs. This Env population is decreased by treatment with BMS-806 or crosslinkers or by the introduction of State-1-stabilizing changes in Env (167). Conversely, this Env population is increased by State-1-destabilizing Env changes that promote CD4 independence (107), by the binding of sCD4 to Env on virus particles, and by CD4 coexpression in virus-producing cells. These cleaved Env trimers apparently represent more open conformations downstream of State 1. The intersubunit interactions in these trimers are more labile than those in the State-1 Envs, as gp120-directed pNAbs less efficiently coprecipitate gp41 in detergent lysates.

### 3) gp41 molecules not detectably associated with gp120

This gp41-only population is marked by its recognition by C34-Ig and the F240 pNAb, which detect gp41 molecules after gp120 has been shed (21,158,186). As seen in our results, the gp41 from this population is precipitated by C34-Ig and F240, but gp120 is not coprecipitated. State-1 stabilization by Env changes or BMS-806 treatment reduces gp120 shedding and decreases the level of gp41-only Envs, implying that the gp41-only population increases under conditions in which more open Env conformations are favored.

**FIG 9.**
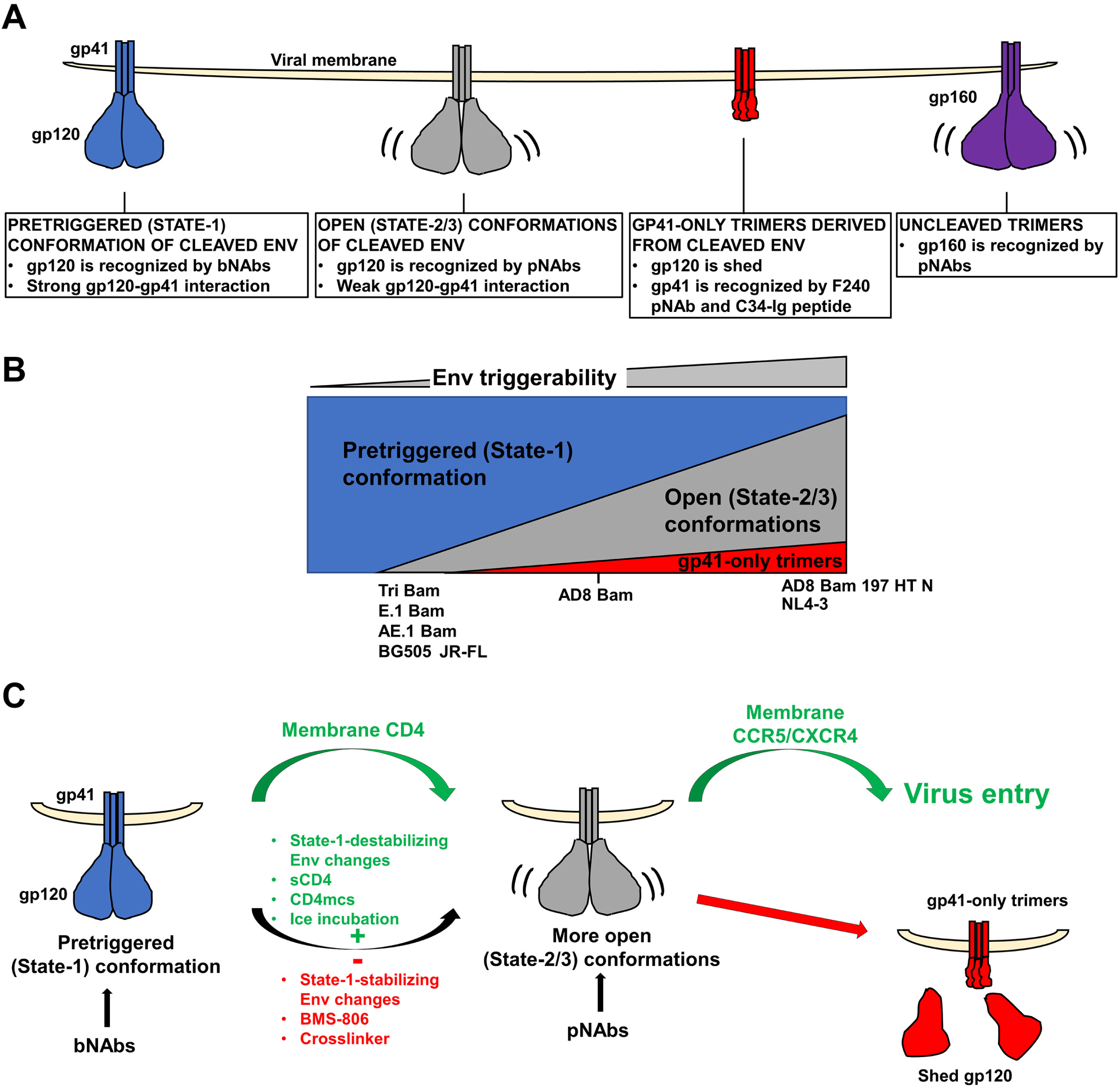
Model of Env conformations on virus particles. (A) Three populations of cleaved Env trimers on virus particles are depicted along with their distinguishing properties. Uncleaved (gp160) Env trimers, which are found to various extents on VLP preparations, are shown on the right. (B) The relative levels of cleaved Env trimer populations on virions are depicted for HIV-1 Envs with different levels of triggerability. The approximate relationship of the different HIV-1 Env variants used in this study is shown. Env triggerability is inversely related to the activation barrier separating State 1 from States 2/3 and varies among Envs from different primary HIV-1 strains (107). Envs with intermediate levels of triggerability, like the AD8 Bam Env, populate the pretriggered (State-1) conformation and also spontaneously sample more open, downstream (States 2/3) conformations. Viral Envs with lower triggerability (e.g., the Tri Bam or E.1 Bam Envs) populate the pretriggered (State-1) conformation on the virions to a greater extent. Conversely, viral Envs with higher triggerability (e.g., the AD8 Bam 197 HT N Env) exhibit more open trimer conformations on the virions and are more prone to shed gp120, leading to gp41-only trimers. (C) Modulation of virion Env conformational transitions. Upon binding to membrane CD4 on a target cell, the pretriggered (State-1) Env conformation undergoes transitions to more open (State-2/3) intermediates in which the gp120 coreceptor-binding site is exposed. Env binding to the CCR5 or CXCR4 coreceptor promotes additional conformational changes in Env that facilitate virus entry (green arrows). Depending on its triggerability, HIV-1 Env will spontaneously sample more open (State-2/3) conformations (curved black arrow) in which epitopes for some pNAbs become exposed. Such spontaneous transitions from State 1 can be suppressed by State-1-stabilizing Env changes or by treatment of the Env trimers with BMS-806 or chemical crosslinkers (red minus sign). Conversely, State-1-destabilizing Env changes or treatment with sCD4 or CD4-mimetic compounds (CD4mcs) drive Env trimers out of State 1 and increase the level of virion Envs in more open conformations (green plus sign). When in close proximity to a potential target cell expressing CCR5 or CXCR4 coreceptors, virions with Envs in these open (State-2/3) conformations can infect the cell (168,169,171,187,188). However, compared with Envs that engage CD4 on a target membrane, Envs opened by other means are more prone to either spontaneous inactivation or neutralization by pNAbs (36,49,187,188,195). The pNAb-reactive, open Env intermediates exhibit weak intersubunit interactions compared to the bNAb-reactive State-1 Env trimers. Certain HIV-1 isolates like HIV-1_AD8_ are not neutralized by pNAbs yet show pNAb binding to cleaved Env on virus particles; this apparent paradox can be resolved if the cleaved Env recognized by pNAbs is partially or completely dysfunctional. Our results suggest that gp120 shedding is more likely to occur from more open Env conformers (red arrow).

Our results support a model in which the triggerability or reactivity of Env variants determines the spontaneous occupancy of the three cleaved Env populations on the virus particles (12,48,49,107,113,167–169) (Fig. 9B). Virions with the tier-2 AD8 Bam Env, with an intermediate level of triggerability (167), contain all three cleaved Env populations in the following order of decreasing amount: pretriggered ≥ open >>> gp41-only. Virions with the more triggerable tier-1 NL4-3 Env and the CD4-independent AD8 Bam 197 HT N Env contain all three populations in the following order of decreasing amount: open > pretriggered >>> gp41-only. Virions with the State-1-stabilized Tri Bam and AE.1 Bam Envs and virions with the tier-2/3 JR-FL and BG505 Envs contain only two populations in the following order of decreasing amount: pretriggered > open. We suggest that the lower triggerability or reactivity of this last group of Env variants is related to the higher activation energy barrier separating the pretriggered (State-1) conformation and downstream, more open conformations, leading to higher occupancy of the pretriggered state on the virus particles (107, 168).

The degree to which the cleaved Env trimers on virus particles sample State 1 or more open conformations is subject to modulation (Fig. 9C). CD4 binding promotes HIV-1 entry by stimulating Env transitions from State 1 to downstream conformations (12). Spontaneous transitions from the pretriggered (State-1) conformation of Env can be inhibited by treatment of the virions with BMS-806 (21), by crosslinking, or by the introduction of State-1-stabilizing changes in Env. Conversely, treatment of the virions with sCD4 or a CD4mc, or the introduction of State-1-destabilizing changes in Env can promote Env transitions from State 1 to more open downstream conformations. These downstream Env conformations are recognized by pNAbs that bind the coreceptor-interactive region of gp120, consistent with their potential relevance to virus entry events following CD4 engagement. We note that crosslinking reduces the binding of these pNAbs to the unliganded AD8 Env (Fig. 2D), suggesting that Env flexibility contributes to the spontaneous exposure of the pNAb epitopes. This is consistent with smFRET observations indicating that State-1 Envs spontaneously and reversibly sample conformations that resemble the States 2 and 3 induced by CD4 (12). If these epitopes are accessible on the unliganded virion Env, why don’t the V3 and CD4i pNAbs neutralize HIV-1_AD8_? The lability of gp120-trimer association observed for the AD8 Env population that spontaneously exposes pNAb epitopes hints that these Envs may be inherently dysfunctional or that their functionality is short-lived. Poorly functional Envs with exposed pNAb epitopes have been previously observed on VLPs and were shown to be more susceptible to digestion with a cocktail of proteases (127). Like the short-lived Env intermediates induced by sCD4 or CD4mcs (187, 188), spontaneously sampled, open Env conformations may be labile. To maintain some infectivity, highly triggerable Envs like AD8 Bam 197 HT N may need to acquire adaptive changes to minimize the lability of downstream Env intermediates during virus infection (107). Of note, the complexes formed by virion Env with CD4 on the target cell membrane were found to be significantly more stable than those formed by sCD4 (187).

Our study provides guidance for efforts to improve VLPs as immunogens that present the membrane-anchored, pretriggered (State-1) conformation of HIV-1 Env to the immune system. The first requirement is that VLPs should contain as little gp160 as possible. Uncleaved Env on VLPs or VLP-producing cells is conformationally flexible and could divert desirable immune responses by presenting immunodominant pNAb epitopes to the immune system. Although cleaved Env is generally enriched in virions, some uncleaved Env is found in most VLP preparations. Previous studies with A549 cells stably producing Gag-mCherry VLPs indicated that essentially all of the uncleaved VLP Env passed through the Golgi apparatus (7); the uncleaved Env on VLPs produced in that system was modified by complex carbohydrates resembling those on mature Env. In VLPs produced transiently from 293T cells in this study, some of the uncleaved Env lacks complex glycans, indicating that it may have bypassed the Golgi apparatus. One likely source of this uncleaved Env may be cellular vesicles that contaminate the VLP preparation. The level of vesicle contaminants could be influenced by the choice of producer cells, transient versus stable transfection and the VLP production system. If complete elimination of the uncleaved gp160 Env in VLP preparations is not possible, BMS-806 and long-acting BMS-806 analogues could be used to reduce the exposure of pNAb epitopes on the residual gp160 (38, 118).

The second requirement is that cleaved Env should be in a State-1 conformation with minimal spontaneous exposure of pNAb epitopes. We found that even cleaved HIV-1 Env trimers of the tier-2 AD8 Env spontaneously exposed V3 and CD4i pNAb epitopes related to the coreceptor-binding region of gp120. The extent of this exposure appears to be related to Env triggerability, i.e., the propensity of Env to make transitions from the pretriggered (State-1) conformation to downstream conformations (107, 168). State-1-stabilizing Env changes, BMS-806 treatment and crosslinking (or combinations of these measures) can decrease the amount of the more open cleaved Envs trimers on the VLPs, even beyond that found in natural HIV-1 strains. As State-1 stabilization improves, Env typically demonstrates decreased ability to mediate virus infection, as expected for lower triggerability (107, 167). Assays dependent on functional virus (e.g., antibody neutralization, cold sensitivity, inhibition by sCD4 or small-molecule entry blockers) cannot be used to study such highly State-1-stabilized Envs. The VLP antigenicity and gp120 shedding assays developed here do not require the virus to be functional, and thus can be applied in broad context to monitor progressive improvements in the stabilization of the pretriggered conformation of Envs from multiple HIV-1 strains. Fortunately, bNAbs and pNAbs broadly reactive with the Envs from diverse HIV-1 are available for the antigenicity analyses, and thus the antigenicity of different HIV-1 strains can be directly compared to allow Env conformations to be deduced.

The third requirement is that shedding of gp120 from the cleaved Env on VLPs should be minimized. Viruses with the AD8 Bam Env spontaneously shed gp120 after incubation at various temperatures (Fig. 5). Trimer stability after incubation of the VLPs on ice is a better indicator of State-1 stabilization than at higher temperatures (167,172,173). The detrimental effects of ice formation at near-freezing temperatures apparently stress the non-covalent intersubunit interactions on which Env trimer integrity depends (189–191). Further studies of the longevity of State-1-stabilized Env trimers at 37°C would be relevant to their inclusion in VLP immunogens, particularly in the presence of adjuvants. Encouragingly, State-1-stabilized Envs like the Tri Bam and AE.1 Bam Envs and the natural JR-FL Env were remarkably resistant to gp120 shedding. Using these Envs in VLPs provides alternatives to the use of artificial inter-subunit disulfide bonds like SOS (192–194), which has been reported to destabilize State 1 in virion Envs (95, 118).

The fourth requirement is that VLP-producing cells should not coexpress CD4. CD4 expression in the VLP-producing cells was found to exert subtle effects on the exposure of pNAb epitopes on the VLP Envs, even for Vpu^+^ Nef^+^ proviruses (Fig. 8B). The use of CD4-negative cells to produce VLPs enriched in State-1 Envs seems advisable.

Advances in understanding the pretriggered (State-1) conformation of HIV-1 Envs and learning how to preserve this labile state, together with the assays established here, should assist efforts to elicit effective antibody responses with VLP and other immunogen formulations.

## MATERIALS AND METHODS

### Plasmids

The HIV-1_AD8_, HIV-1_AE.2_ and HIV-1_JR-FL_ *env* sequences for the construction of IMCs were obtained from the respective pSVIIIenv expression vectors (167). Relative to the AD8 Env, the AE.2 Env contains the following changes: Q114E, R166K, R178K, R252K, R315K, R419K, R557K, Q567K, A582T, R633K, Q658K, A667K and N677K. The Kpn I – BamHI *env* fragments from pSVIIIenv AD8 and pSVIIIenv AE.2 were introduced into the pNL4-3 IMC, using an intermediary vector, pE7SB-NL4-3. The pE7SB-NL4-3 plasmid contains the Sal I – BamHI fragment of the HIV-1_NL4-3_ provirus, which includes the *tat*, *rev*, *vpu* and 5’ *env* genes. The Kpn I – BamHI *env* fragments from the pSVIIIenv plasmids were cloned into the corresponding sites of pE7SB-NL4-3 using Long Ligase (Takara) following the manufacturer’s protocol. The Sal I – BamHI fragments from the pE7SB-NL4-3 intermediate plasmids were cloned into the corresponding sites of pNL4-3, which contains the infectious HIV-1_NL4-3_ provirus (NIH HIV Reagent Program). The resulting pNL4-3.AD8 and pNL4-3.AE.2 IMCs express AD8 and AE.2 Envs with N-terminal residues 1-33 (including the signal peptide) and C-terminal residues 751-856 (C-terminus of Env cytoplasmic tail) from the NL4-3 Env.

The pNL4-3.AD8 Bam and pNL4-3.AE.2 Bam IMC was created by introducing mutations encoding S752F and I756F changes in the Env cytoplasmic tail into pNL4-3.AD8 and pNL4-3.AE.2 using primers forward: tccgtgcgattagtggatggatTcttggcacttTtctgggacgatctgcggagcctgtgcctcttca, reverse: tctgtctctgtctctgtctccaccttcttcttcgattccttcgggcctgtcgggtcccctcggggct. The pNL4-3.AD8(-) IMC encodes an AD8 Env in which the REKR cleavage site (residues 508-511) is altered to SEKS. The pNL4-3.AD8 Δ712 plasmid encodes an AD8 Env with a truncated cytoplasmic tail (missing residues 712-856).

To knock out HIV-1 genes in the pNL4-3.AD8 plasmid, stop codons were introduced individually into the open reading frames encoding RT, RNAse H, IN, Vif, Vpr, Vpu and Nef. The primers for these knockouts were: RT forward: TTTAAATTTTtaaATTAGTCCTATTGAGACTGTAC, reverse: GTGCAGCCAATCTGAGTC; RNAse H forward: AAACTTTCTAaGTAGATGGGGC, reverse: CTGCTCCTATTATGGGTTC; IN forward: AAAGTACTATaaTTAGATGGAATAGATAAGGC, reverse: CCTGATTCCAGCACTGAC; Vif forward tgaggattaacacaTAGaaaagattagtaaaa, reverse tcctgtctacttgccacacaatcatcacctgc; Vpr forward: actgacagaggacagatggaaTAAgccccagaagaccaa, reverse: ttcctaacactaggcaaaggtggctttatctgtt; Vpu forward: TAgcaacctataatagtagcaatagtagcattagtagtagca reverse: tacatgtactacttactgctttgatagagaagcttgat Nef forward: TAAggtggcaagtggtcaaaaagtagtgtga reverse: cttatagcaaaatcctttccaagccctgtctt.

IMCs encoding the Tri Bam and AD8 Bam 197 HT N Envs were made by introducing Q114E + Q567K + A682T or N197S + NM 625/626 HT + D674N changes, respectively, into pNL4-3.AD8 Bam IMC.

The pNL4-3.AE.1 Bam IMC was cloned from pNL4-3.AE.2 Bam by back reverting the following lysine residues: K658Q, K667A and K677N. The pNL4-3.E.1 Bam was cloned by back reverting T582A in pNL4-3.AE.1 Bam. The Strep tag was inserted at the C terminus by site-directed mutagenesis using primers forward: CCCAGTTCGAGAAAtaagatgggtggcaagtggtcaaaaagtagtgtgat, reverse: GGTGGCTCCAtagcaaaatcctttccaagccctgtcttattcttctaggta.

To create IMCs expressing soluble gp120s, the codon for gp120 residue 508 was replaced by a stop codon. All site-directed mutagenesis with IMCs was done using the Q5 high-fidelity DNA polymerase (New England Biolabs) and One Shot Stbl3 Chemically Competent E. coli (Invitrogen) following the manufacturer’s protocols.

To clone the pNL4-3.JR-FL IMC, an overlap extension PCR using PfuUltra II fusion HS DNA Polymerase (Agilent) was performed. The primers and template plasmids are as follows: AD8 Sal I forward (CAACAACTGCTGTTTATCC) and AD8/JRFL Env reverse (ctgtagcactacagatcatc) using the pNL4-3.AD8 template; AD8/JRFL Env forward (gatgatctgtagtgctacag) and JRFL BamHI reverse (gtcccagataagtgccaag) using the pSVIIIenv JR-FL template (167). The resulting Sal I – BamHI *vpu*_NL4-3_-*env*_JR-FL_ fragment was transferred to the pNL4-3.AD8 vector using Long Ligase.

To clone the pNL4-3.BG505 IMC, an overlap extension PCR was performed. The primers and template plasmids are as follows: pNL4-3.AD8 Sal I forward (CAACAACTGCTGTTTATCCATTTCAGAATTG) and BG505 vpu reverse (AATTTCCAAAGGAAGCATtacatgtactacttactg) using the pNL4-3.AD8 template; BG505 vpu forward (cagtaagtagtacatgtaATGCTTCCTTTGGAAATT) and pcDNA BG505 BamHI mut reverse (GCAAGAGCTAAGgATCCGCTCAC) using the pcDNA3.1 BG505 template (NIH HIV Reagent Program). The Sal I – BamHI *vpu*_BG505_-*env*_BG505_ fragment was transferred to the pNL4-3.AD8 vector using Long Ligase.

### Antibodies and sCD4

The following reagents were obtained through the NIH HIV Reagent Program, Division of AIDS, NIAID, NIH: VRC03, PGT121, 4E10, 10E8.v4, 39F, 3074, 3869, 17b, E51 and sCD4.

### Cell lines

HEK 293T, HeLa cells and TZM-bl cells (ATCC) were cultured in Dulbecco’s modified Eagle’s medium (DMEM) supplemented with 10% fetal bovine serum (FBS) and 100 μg/ml penicillin-streptomycin (Life Technologies). CCR5-expressing C8166-R5 T cells were cultured in Roswell Park Memorial Institute (RPMI) 1640 medium supplemented with 10% FBS and 100 μg/ml penicillin-streptomycin; 1 μg/ml of puromycin was added every fifth passage.

### Env expression and incorporation into virus particles

To prepare pseudoviruses, 293T cells were cotransfected using polyethyleneimine (PEI, Polysciences) with an Env-expressor plasmid, a Tat-encoding plasmid and the pNL4-3.ΔEnv plasmid (renamed from pNL4-3.Luc.R-E-plasmid, available at the NIH HIV Reagent Program) at a 1:0.125:1 weight ratio unless indicated otherwise. To prepare replication-competent viruses, 293T cells were transfected with the pNL4-3.Env plasmid using PEI. The medium was replaced at 4-6 h after transfection. Lipofectamine 3000 (Invitrogen) was used to transfect HeLa cells following the manufacturer’s protocol. At 48 h to 72 h after transfection, cells were lysed and clarified; supernatants were collected, filtered (0.45 μm) and pelleted at 14,000-100,000 x g for 1 h at 4°C. Virus pellets and clarified cell lysates were then analyzed by Western blotting using a nitrocellulose membrane and wet transfer (350 A, 75 min, Bio-Rad). Western blots were developed with 1:2,000 goat anti-gp120 polyclonal antibody (Invitrogen), 1:2,000 4E10 anti-gp41 antibody, 1:1,000 mouse anti-p24 serum (John C. Kappes, University of Alabama at Birmingham), 1:10,000 rabbit anti-hsp70 (K-20) antibody (Santa Cruz Biotechnology). The HRP-conjugated secondary antibodies were 1:2,000 rabbit anti-goat (Invitrogen), 1:2,000 goat anti-human (Invitrogen), 1:1,000 goat anti-mouse (Invitrogen) and 1:10,000 goat anti-rabbit (Sigma-Aldrich). The intensity of protein bands on non-saturated Western blots was quantified using the Bio-Rad Image Lab program. Statistical significance was evaluated by a two-tailed Student’s t test.

### Virus infectivity

The cell supernatant containing virus was clarified by low-speed centrifugation (2,000 rpm for 10 min). To compare the infectivity of different viruses, an equal volume of clarified supernatant was incubated with TZM-bl cells in 96-well plates (2 x 10^4^ cells per well). The plates were incubated at 37°C/5% CO_2_ for 48 h, after which the cells were lysed and luciferase activity was measured using a luminometer.

### Virus neutralization

Approximately 100 to 200 TCID_50_ (50% tissue culture infectious dose) of virus was incubated with serial dilutions of purified antibodies, sCD4-Ig or BNM-III-170 at 37°C for 1 h. The mixture was then added to TZM-bl cells in 96-well plates and luciferase activity was measured after 48 h as described above. The concentrations of antibodies and other inhibitors that inhibit 50% of infection (the IC_50_ values) were determined using GraphPad Prism 8 (five-parameter dose-response) or Microsoft Excel graphs.

### Deglycosylation of Env on virus particles

Purified virus particles were first lysed in 1X PBS/0.5% NP-40. The virus lysate was then denatured by boiling in denaturing buffer (New England BioLabs) for 10 min and treated with PNGase F or Endo Hf enzymes (New England BioLabs) for 1.5 h at 37°C in accordance with the manufacturer’s protocol. The treated proteins were then analyzed by reducing SDS-PAGE and Western blotting.

### Antigenicity of Env on virus particles

To assess Env antigenicity, 50-100-µl aliquots of virus particles (purified and resuspended in 1X PBS) were incubated with a panel of antibodies at 10 µg/mL concentration for 1 h at room temperature. One mL of chilled 1X PBS was added and samples were centrifuged at 14,000 x g for 1 h at 4°C. The pellets were lysed in 100 µl chilled 1X PBS/0.5% NP-40/protease inhibitors cocktail. VLP lysates were rotated during incubation with Protein A-agarose beads for 1 h at 4°C and washed with chilled 1X PBS/0.1% NP-40 three times. The beads were resuspended in 1X PBS containing NuPage LDS Sample Buffer (New England Biolabs) and dithiothreitol (DTT) and used for Western blotting. To prepare the Input (50%) sample, half of the virus volume was mixed with 1 mL chilled 1X PBS and centrifuged at 14,000 x g for 1 h at 4°C; the pellet was resuspended in 1X PBS/LDS/DTT. To examine the effects of soluble CD4 or BMS-806 on Env conformation, 10 µg/mL four-domain sCD4 or 10 µM BMS-806 was first added to virus particles before antibodies were added. To examine the antigenicity of crosslinked Env, a concentrated volume of virus particles was first incubated with 0.1 mM or 1 mM DTSSP crosslinker for 30 min at room temperature, after which the reaction was quenched with 100 mM Tris-HCl, pH 8.0, for 10 min at room temperature. More PBS was added and Env antigenicity was evaluated as described above.

### Oligomerization of Env on virus particles

Purified virus particles were incubated with different concentrations of BS3 crosslinkers (ThermoFisher Scientific) for 30 min at room temperature, after which the reaction was quenched with 100 mM Tris-HCl, pH 8.0, for 10 min at room temperature. LDS/DTT was added and samples were boiled and then analyzed by reducing SDS-PAGE and Western blotting.

### Antigenicity of soluble gp120

293T cells were transfected with IMCs expressing the soluble gp120 version of the AD8 Bam, Tri Bam, AE.1 Bam, NL4-3, JR-FL or BG505 Envs using PEI. Forty-eight hours after transfection, cell supernatants containing the soluble glycoproteins were collected and filtered (0.45 mm). Aliquots were incubated with 10 µg/mL antibody and Protein A-agarose beads, and the mixture was rotated at room temperature for 2 h. The beads were washed three times with 1X PBS/0.1% NP-40 before the beads were boiled and Western blotted with a goat anti-gp120 antibody, as described above. To crosslink soluble gp120, the filtered supernatants from transfected 293T cells were incubated with 1 mM DTSSP for 30 min at room temperature. The reactions were quenched with 100 mM Tris-HCl, pH 8.0, for 10 min before the antigenicity assay was carried out as described above.

### Shedding of virus particles

Purified virus particles were resuspended in 1X PBS, aliquoted into 50 μL and incubated with serial dilutions of sCD4 or the CD4mc BNM-III-170 for 1 h at room temperature or the indicated temperatures. Next, 200 μL 1X PBS was added and samples were centrifuged at 14,000 x g for 1 h at 4°C. Then 220 μL of the supernatants was collected and rotated during incubation with Galanthus Nivalis Lectin (GNL)-agarose beads (Vector Laboratories) for 2 h at room temperature. Beads were washed three times with 1X PBS/0.1% NP-40 and processed for Western blotting with a goat anti-gp120 antibody, as described above. To evaluate spontaneous gp120 shedding at different temperatures, aliquots of purified virus particles were incubated on ice or at different temperatures for the indicated amount of time before the samples were processed as described above. An aliquot of the purified virus particles prior to incubation with ligands or at different temperatures was used as the “Input” sample.

### Antigenicity of Env virions from infected T cells

293T cells were transfected with IMCs expressing the Strep-tagged AD8 Bam or E.1 Bam Env using PEI. Seventy-two hours after transfection, the cell supernatants were clarified by low-speed centrifugation and serial dilutions of the virus were used to infect TZM-bl cells. Forty-eight hours later, the luciferase activity in the TZM-bl lysates was measured. Virus TCID_50_ values were calculated using the Reed-Muench method (196, 197). Approximately 10^7^ C8166-R5 cells at a density of 1 x 10^6^ cells/mL were infected with virus at a multiplicity of infection of 0.1 for 5 h or overnight, in the presence of 8 μg/mL polybrene. The cells were then washed and fresh medium was added to dilute the cells to a density of 2.5 x 10^5^ cells/mL. At three days after infection, one-quarter of the cells/medium was saved and diluted 1:4 with fresh medium. Six to seven days after infection, virus particles were collected, purified and concentrated, and Env antigenicity was analyzed as described.

## ACKNOWLEDGMENTS

We thank Ms. Elizabeth Carpelan for manuscript preparation. Antibodies against HIV-1 were kindly supplied by John C. Kappes (University of Alabama at Birmingham), Dennis Burton (Scripps), Peter Kwong and John Mascola (Vaccine Research Center NIH), Barton Haynes (Duke University), Hermann Katinger (Polymun), James Robinson (Tulane University), and Marshall Posner (Mount Sinai Medical Center). We thank the NIH HIV Reagent Program for providing additional reagents.

This work was supported by grants from the National Institutes of Health (grant nos. AI 145547, AI 124982, AI 150471, AI 129017 and AI 164562), a grant from Gilead Sciences, and by a gift from the late William F. McCarty-Cooper.

We declare no conflicts of interest.

